# Reading frame restoration at the EYS locus, and allele-specific chromosome removal after Cas9 cleavage in human embryos

**DOI:** 10.1101/2020.06.17.149237

**Authors:** Michael V. Zuccaro, Jia Xu, Carl Mitchell, Diego Marin, Raymond Zimmerman, Bhavini Rana, Everett Weinstein, Rebeca T. King, Morgan Smith, Stephen H. Tsang, Robin Goland, Maria Jasin, Rogerio Lobo, Nathan Treff, Dieter Egli

**Affiliations:** Division of Molecular Genetics, Department of Pediatrics and Naomi Berrie Diabetes Center, Columbia University, New York, NY 10032, USA; Columbia Stem Cell Initiative, Columbia University, New York, NY 10032, USA; Genomic Prediction Inc. 675 US Highway One, Suite 126 North Brunswick, NJ 08902, USA4^3^ Division of Reproductive Endocrinology and Infertility, Department of Obstetrics and Gynecology, Columbia University, New York, NY 10032, USA; Department of Molecular Biosciences, Rutgers, The State University of New Jersey, New Brunswick, NJ 08854, USA; Jonas Children’s Vision Care, and Bernard & Shirlee Brown Glaucoma Laboratory, Departments of Ophthalmology, Pathology & Cell Biology, Institute of Human Nutrition, Vagelos College of Physicians and Surgeons, Columbia University. New York, New York, 10032, USA; Developmental Biology Program, Memorial Sloan Kettering Cancer Center, New York, NY, USA; Division of Reproductive Endocrinology and Infertility, Department of Obstetrics and Gynecology, Columbia University, New York, NY 10032, USA

**Author notes:** equal contribution (Michael V. Zuccaro, Jia Xu).

**Keywords:** Human embryo, Cas9, gene editing, germ line, chromosome loss, double strand break repair, homologous recombination, microhomology mediated end joining, mitosis, interhomolog recombination

## Abstract

The correction of disease-causing mutations in human embryos could reduce the burden of inherited genetic disorders in the fetus and newborn, and improve the efficiency of fertility treatments for couples with disease-causing mutations in lieu of embryo selection. Here we evaluate the repair outcomes of a Cas9-induced double-strand break (DSB) introduced on the paternal chromosome at the *EYS* locus, which carries a frame-shift mutation causing blindness. We show that the most common repair outcome is microhomology-mediated end joining, which occurs during the first cell cycle in the zygote, leading to embryos with non-mosaic restoration of the reading frame. However, about half of the breaks remain unrepaired, resulting in an undetectable paternal allele and, upon entry into mitosis, loss of one or both chromosomal arms. Thus, Cas9 allows for the modification of chromosomal content in human embryos in a targeted manner, which may be useful for the prevention of trisomies.

**Highlights:**

- Cas9-mediated DSB induction and repair by end joining occurs within hours
- End joining provides an efficient way to restore reading frames without mosaicism
- Unrepaired DSBs persist through mitosis and result in frequent chromosome loss

## Background

Double-strand breaks (DSBs) stimulate recombination between homologous DNA segments (Jasin and Rothstein, 2013). The targeted introduction of a DSB followed by recombination allows for the precise modification of genomes in model organisms and cell lines, and may also be useful for the correction of disease-causing mutations in the human germ line (Lea and Niakan, 2019). DSBs occur naturally during meiosis, and are repaired through recombination between homologous chromosomes, thereby ensuring genome transmission and genetic diversity in offspring. Recombination between homologs was also recently suggested to occur efficiently in mitotically dividing cells: a DSB at the site of a disease-causing mutation on the paternal chromosome resulted in the loss of the mutation such that approximately half of the resulting embryos carried only the maternal wild-type allele (Ma et al., 2018; Ma et al., 2017). The elimination was presumed to occur through use of the maternal genome as a repair template, resulting in what appeared to be the efficient correction of a pathogenic mutation on the paternal chromosome without mosaicism. This contrasts with frequent mosaicism in previous studies with different cells of the same embryo carrying various edited and non-edited alleles (Liang et al., 2015).

The correction of pathogenic mutations through interhomolog recombination with a lack of mosaicism, if independently confirmed, would have major advantages over other approaches, since such a mechanism of correction does not require the introduction of exogenous nucleic acids and is limited to alleles already present in the human population. However, alternative interpretations of the results have been proposed, including the loss of the paternal allele through large deletions, chromosome loss, or translocations (Adikusuma et al., 2018; Egli et al., 2018). Furthermore, the physical location of the genomes during the first cell cycle pose a significant limitation to the occurrence of such a recombination event, as maternal and paternal genomes are packaged in separate nuclei and only come together in the same nucleus after the first mitosis at the two-cell stage (Reichmann et al., 2018). Thus, the timing of Cas9-induced breakage and repair is an important determinant of the genetic outcomes with regard to mosaicism and the mechanisms available for repair. Many questions remain regarding DSB repair in human embryos, in particular because of a limited ability to detect complex repair events in a small number of cells.

Here we use sperm from a patient with blindness due to a homozygous frame-shift mutation in the *EYS* gene located on chromosome 6 to analyze the timing and the mechanisms of double-strand break (DSB) repair in human embryos. We show that a Cas9-induced break on the paternal genome within the first cell cycle post fertilization is repaired through nonhomologous end joining (NHEJ) and microhomology-mediated end joining (MMEJ) in half of the embryos. In the majority of end joining events, the reading frame of the *EYS* gene is restored, allowing uniform nonmosaic embryo editing. Although the other half of the embryos appear as wild type in an on-target genotyping analysis, they arise not by correction, but instead through both segmental and whole chromosome loss due to mitotic entry with unrepaired breaks. Though these embryos develop to the expanded blastocyst stage, they fail to establish embryonic stem cell lines. These results show a surprising tolerance of the human embryo for cell cycle progression despite unrepaired DSBs, and suggest that targeted pericentromeric cleavage by Cas9 may enable the allele-specific removal of chromosomes for the *in vitro* correction of trisomies.

## Results

### Allele-specific editing of the *EYS* gene in embryonic stem cells

We recruited a sperm donor with a homozygous G deletion mutation (rs758109813) in the gene encoding *EYS* associated with retinitis pigmentosa (Abd El-Aziz et al., 2008), which results in the frame-shift p.Pro2265Glnfs*46 (referred to as *EYS^2265fs^)* in exon 34 (**Fig. 1A**). The *EYS* gene is located on chromosome 6, at 6q12, near the centromere of the long arm. A guide RNA (gRNA) was designed to specifically target the mutant but not the wild-type allele, which differs at the PAM sequence motif. The Cas9 cleavage site occurs proximal to *EYS^2265fs^*, such that small indels will preserve the original SNP, which can distinguish modified alleles of paternal or maternal origin. Furthermore, the mutation is flanked by common SNPs rs66502009, centromeric to the mutation, rs12205397 and rs4530841, both telomeric to the mutation, which are amplified within a single ~1kb PCR product, as well as by SNPs at greater distance. Thus, oocytes from donors with homozygous SNPs that differ from the sperm donor allow the identification of maternal and paternal chromosomes in the embryo, and the evaluation of novel combinations of maternal and paternal alleles (**Fig. 1A**, **Table S1**).

**Figure 1:**
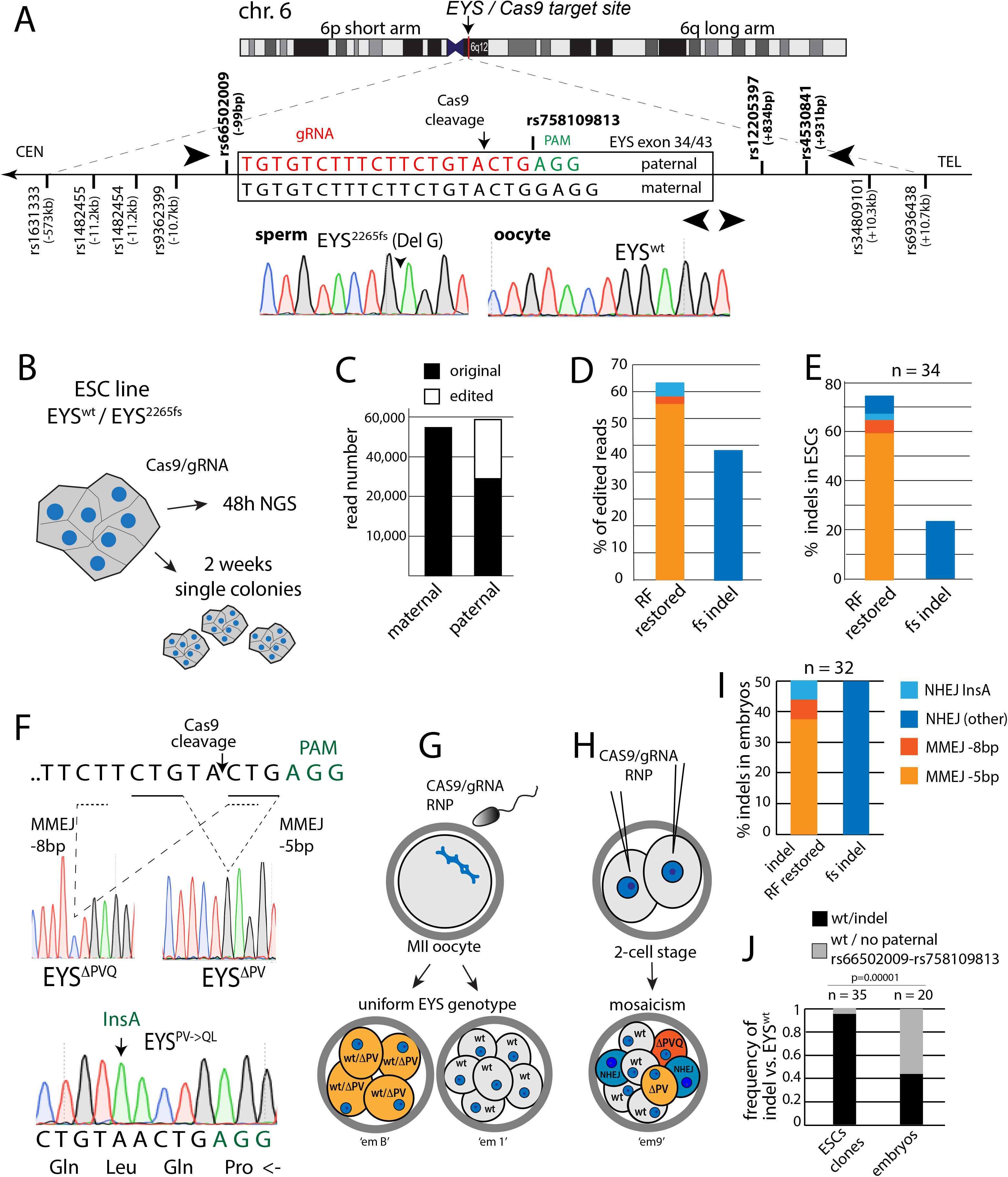
Efficient end joining within the first cell cycle after Cas9 RNP injection at fertilization. **A)** Schematic of genotypes at the human EYS locus with the paternal homozygous EYS^2265fs^ mutation. Alignment of paternal and maternal alleles, gRNA target and flanking SNPs are indicated. Flanking primers (top arrowheads) were used for amplification and sequencing of the mutation site and linked SNPs within a single PCR product. Internal primers (bottom arrowheads) were used for Sanger sequencing and on-target NGS. CEN=centromere, TEL=telomere. Direction of *EYS* transcription is towards the centromere, indicated by the arrow. Sanger sequences of oocyte and sperm donor with homozygous different flanking SNPs. **B)** Schematic of gRNA specificity testing in embryonic stem cells (ESC) with the same genotypes as the fertilized zygote. 48h after Cas9-GFP nucleofection, cells are harvested and used for on-target NGS of the mutation site and rs66502009. **C)** Read quantification of edited and original alleles. **D**) Type and frequency of indels in human pluripotent stem cells evaluated using either on-target NGS (**D**) or colony picking and Sanger sequencing (**E**). **F)** Schematic of DSB repair events after Cas9 cleavage. Cas9 cleaves between two regions of microhomology. Alternate products of microhomology-mediated end joining (MMEJ) obtained in human embryos are shown below. Nonhomologous end joining can result in reading frame restoration due to insertion of an A due to the Cas9 overhang. **G)** Schematic of editing outcomes when mutant sperm is injected into the cytoplasm together with a gRNA and RNP complex of Cas9 at the MII stage. **H)** Schematic of the injection performed after fertilization at the 2-cell stage. **I)** Quantification of type and frequency of indels of combined data from MII and 2-cell stage injections. **J)** Frequency of ESC clones or embryos with heterozygous indels versus clones or embryos with loss of paternal alleles and an EYS^wt^ genotype. Statistical analysis was performed using Fisher’s exact test. RNP=ribonucleoprotein.

We first derived an ESC line (eysESC1) through fertilization to determine the specificity of the gRNA for the mutant allele (**Fig. 1B**). Heterozygous ESCs for both the mutation site (*wt/EYS^2265fs^)* and rs66502009 were transfected with Cas9-GFP and gRNA expression vectors to target the *EYS^2265fs^* allele, and GFP positive cells were harvested by flow cytometry 48h post transfection. Using PCR and on-target next-generation sequencing (NGS) of two independent biological samples scoring for SNPs rs66502009 and *EYS^2265fs^*, we found that 48.6% (37,646 of 77,395) were unmodified reads of the maternal *EYS^wt^* allele, while 51.3% (39,749 of 77,395) were of paternal origin, of which 38.5% (15,307 of 39,749) were edited to contain small indels on reads containing both the *EYS^2265fs^* as well as the rs66502009 paternal allele (**Fig. 1C**, **Table S2**). This demonstrates the specificity of the gRNA for the paternal allele in ES cells, resulting in efficient modification at *EYS^2265fs^* (**Table S2**).

### End joining restores the EYS reading frame by inducing predictable indels in ESCs and embryos

Analysis of the types of edits in NGS reads showed that the most frequent event (55% of edited reads) was the deletion of 5bp on the paternal allele (**Fig. 1D**), resulting in the restoration of the reading frame. Deletion of 8bp and insertion of 1bp, both of which restore the reading frame, were also observed. In addition to analysis of NGS reads, we also plated single cells for colony formation representing biologically independent editing events. In 245 clones grown from single cells, we identified 35 edited clones (14.3%), 34 of which had indels in the paternal allele, while one had no detectable paternal allele at either rs66502009 or *EYS^2265^*. Of the 34 clones with indels, the reading frame was restored in 27 (79.4%), primarily through a recurrent 5bp deletion (**Fig. 1E**). The placement of the guide RNA resulted in cleavage between two identical regions of either 3bp or 2bp, defining sites of micro-homology (**Fig. 1F**). Microhomology-mediated end joining (MMEJ) is an efficient repair pathway that uses one to a dozen base pairs of homology (Sfeir and Symington, 2015). MMEJ at the *EYS* locus results in the deletion of either 5bp or 8bp, thereby restoring the reading frame, and generating two different novel alleles with deletions of either two amino acids (p.P2265-V2266) (EYS^ΔPV^) or three amino acids *(p.Pro2265_Gln2267del)* (EYS^ΔPVQ^). The deletion EYS^ΔPV^ was by far the most common single repair product in ESCs. No mitotic recombination between rs758109813 and rs66502009, which would be indicative of interhomolog repair, was observed in the 34 edited clones.

To determine editing outcomes in human embryos, we injected *EYS^2265fs^* mutant sperm into the cytoplasm (ICSI) together with a ribonucleoprotein (RNP) complex of Cas9 nuclease and the gRNA targeting *EYS^2265fs^* (**Fig. 1G**). Alternatively, Cas9/RNP was injected after fertilization at the 2-cell stage (**Fig. 1H**). In both types of injections, embryos were biopsied for genotyping at the cleavage or the blastocyst stage. While injection at the two-cell stage invariably resulted in mosaic embryos (n=13) with up to three genotypes (**Table S1**), injection at the MII stage resulted in embryos (n=7) that appeared uniform, for either an indel (n=3), or only the *EYS^wt^* allele (n=4). Combining both data from MII injections as well as 2-cell stage injections, we found a total of 32 end joining events that were independent, because they occurred in different embryos, or differed molecularly within the same embryo (**Fig. 1I**, **Table S1**). 14 of these were MMEJ events, 12 of which resulted in a 5bp deletion and 2 in an 8bp deletion. Furthermore, of a total of 18 independent NHEJ events, 2 restored the reading frame through insertion of an A nucleotide, resulting in a transition to *(p.Pro2265_Val2266insGlnLeu)* EYS^PV>QL^. Importantly, in one embryo derived from an MII injection, all blastomeres (4/4) showed uniform reading frame restoration, demonstrating nonmosaic editing through MMEJ (**Table S1**), The reproducible and frequent generation of the EYS^ΔPV^ allele in both embryos and stem cells shows that MMEJ can efficiently be used for the restoration of the reading frame at EYS^2265fs^.

Though the frequency of different end joining events in pluripotent stem cells and embryos was comparable, there was one significant difference in editing outcomes: considering both MII and 2-cell stage injections, 17 of 20 embryos contained cells with only an *EYS^wt^* allele and the flanking maternal rs66502009 allele, representing at least 17 independent events, while 12 embryos contained cells with indels. Therefore, the loss of the paternal allele was as or more common than a heterozygous indel in embryos (**Fig. 1J**). Only a single event (2.8%) was seen in Cas9 treated ESC clones. These events could represent unrepaired paternal alleles, undetectably modified paternal alleles (Adikusuma et al., 2018; Kosicki et al., 2018), or interhomolog repair as previously suggested (Ma et al., 2017).

### End joining within the first cell cycle after Cas9 RNP injection at fertilization

To better understand the origin of *EYS^wt^* events, as well as to evaluate the potential for mosaicism in MII injections, we turned to analysis of DNA repair products in zygotes within the first cell cycle. The conclusive determination of nonmosaic embryos requires successful analysis of all cells, which is more reliably achieved when there is just a single genome.

To evaluate the timing of Cas9 cleavage and repair in embryos, we first interrogated repair products in the absence of the maternal genome using androgenesis. The maternal genome was removed and the *EYS^2265fs^* mutant sperm was injected into the cytoplasm (ICSI) together with a ribonucleoprotein (RNP) complex of Cas9 nuclease and the gRNA targeting the mutation site (**Fig. 2A**). At 20h post ICSI, embryos with a single paternal pronucleus were harvested for analysis. Of 8 androgenetic zygotes from two different oocyte donors (5 and 3 oocytes per donor), 3 contained a single small indel, while 2 contained two different modified alleles (**Fig. 2A**, **Table S1**). Of the remaining 3, one was unmodified, and two failed to genotype. Therefore, DSB repair occurred both before and after replication of the *EYS* locus in the androgenetic zygotes.

**Figure 2:**
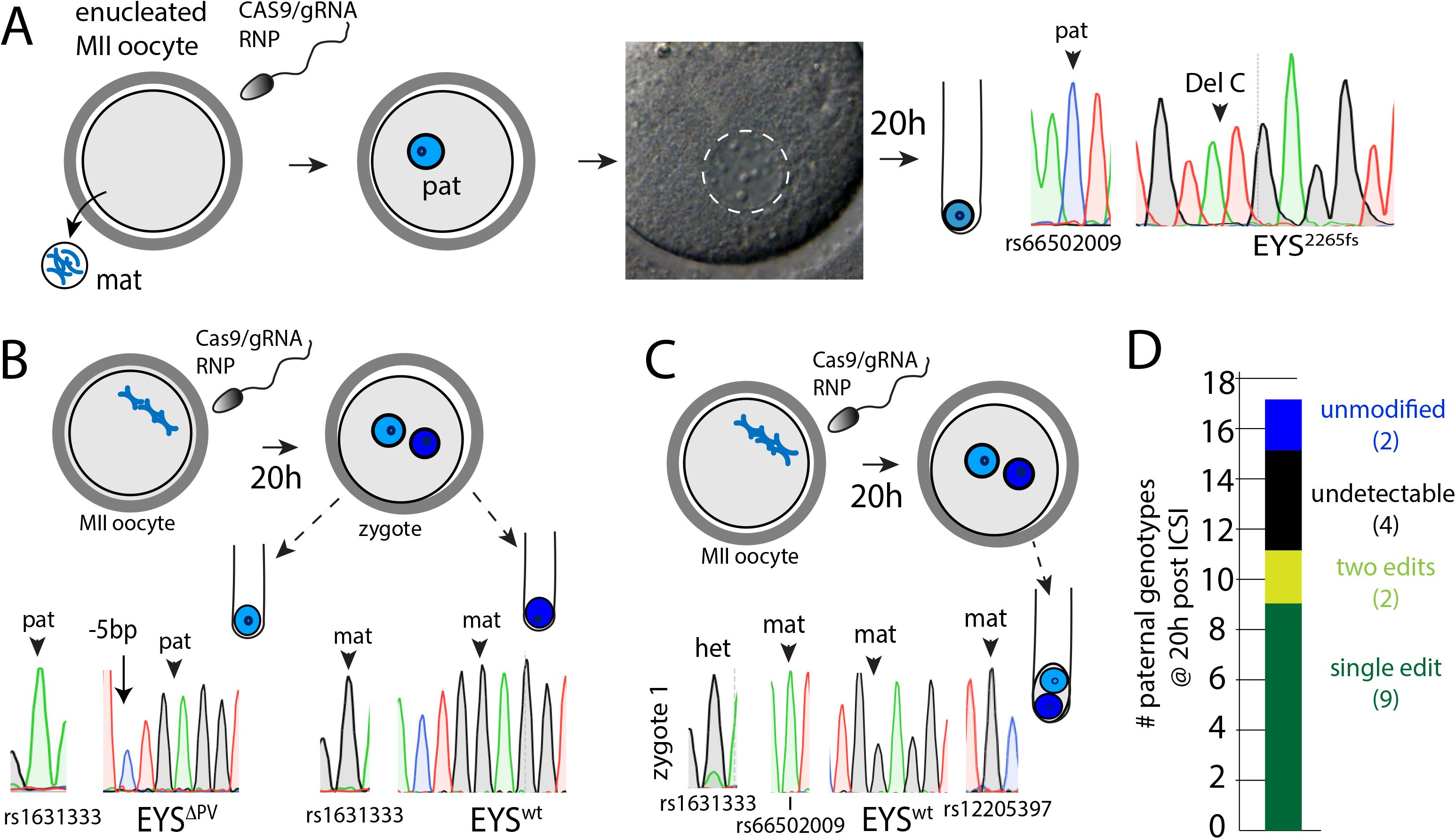
DSB repair occurs during the first cell cycle and independent of the maternal genome. **A**) Evaluating DNA repair outcomes in the absence of the maternal genome. A single sperm together with Cas9 RNP is injected into an enucleated oocyte, resulting in a 1PN zygote, followed by collection at 20h in the first cell cycle for genotyping. **B)** Schematic of fertilization resulting in a zygote with both maternal and paternal genomes. Nuclei isolated and separated to two different tubes for whole genome amplification and genotyping. **C**) ‘Standard’ genotyping analysis where both nuclei of the zygote are collected within the same tube for genotyping. **D**) Summary of genotyping results of the paternal EYS locus at the 1-cell stage.

To determine whether the maternal genome can provide a repair template for the paternal genome and give rise to *EYS^wt/wt^* embryos, we next performed ICSI with Cas9 RNP in nucleated oocytes to generate zygotes with both genomes. To avoid the possibility of allele-dropout due to unequal amplification of alleles, we isolated the individual paternal and maternal nuclei from zygotes at 20h post ICSI and amplified them separately (**Fig. 2B**). Of the 8 zygotes, 5 contained an indel on the paternal allele, while one was unmodified, and two failed to genotype at the mutation site but not at flanking SNPs 10kb away (**Fig. 2B**, **Table S1**). All 5 genomes with indels showed only a single modification. No zygote contained a paternal genome with a wild-type *EYS* allele, and thus there was no evidence for repair from the maternal genome among these zygotes.

To determine the accuracy of genotyping in zygotes with both alleles, i.e. without physical separation of nuclei, we first amplified single cells from embryos (n=3) and single ESCs (n=4) containing both genomes without exposure to the Cas9 RNP. All cells showed heterozygosity at the mutation site as well as at the flanking SNP rs66502009, demonstrating that both alleles were reliably detected (**Table S1**). We next analyzed fertilized zygotes at 20h post fertilization and Cas9 RNP injection (**Fig. 2C**). Of three fertilized oocytes, one zygote showed an edited paternal allele, while two showed only maternal alleles at the mutation site as well as at flanking SNPs within a ~22 kb window on either side (**Fig. 2C**, **Table S1**). For one of the zygotes, a SNP 573kb away could be determined and was found heterozygous for the paternal and maternal alleles (**Fig. 2C**). To account for the possibility of allele dropout in Sanger sequencing, on-target NGS of both ‘wild-type’ zygotes was performed, which showed no significant reads of either the paternal mutation or the flanking paternal SNP at rs66502009 (**Table S1, Table S2**).

Considering all 17 embryos successfully analyzed at the 1-cell stage (20h post ICSI & Cas9 RNP), 11 of the paternal genomes were detectably modified by end joining during this first cell cycle, (**Fig. 2D**, **Table S1**). Editing predominantly gave rise to a single modification (9 out of 11 events) and thus nonmosaic zygotes, whereas 2 of the androgenetic embryos were mosaic. Though these mosaics were seen in androgenetic zygotes, they show that MII injection results predominantly, but not exclusively, in uniform editing. Of the remaining 6 embryos, 2 were unmodified. Four embryos had no detectable paternal *EYS* allele, which had been cleaved and altered by Cas9, although the type of modification is not known. Taken together, 15/17 (88%) paternal alleles were modified by Cas9 in the first cell cycle prior to entry into the first mitosis, including 4 which could not be genotyped.

### Loss of paternal alleles in embryos but not derived stem cell lines

To better characterize the outcomes of a Cas9-induced DSB in zygotes and the developmental potential of edited zygotes, we allowed them to develop to the blastocyst stages for biopsy, genotyping, stem cell derivation, and karyotyping (**Fig. 3A**). Of 18 oocytes injected with sperm and Cas9 RNP at the MII stage, 10 developed to the blastocyst stage (56%). The frequency of blastulation was within the normal range, consistent with earlier reports (Fogarty et al., 2017; Ma et al., 2017). At the cleavage stage, we biopsied blastomeres from 4 embryos, and at the blastocyst stage, we obtained trophectoderm (TE) biopsies consisting of 5-10 cells from 3 embryos (**Fig. 3B**). At the cleavage stage, 2 embryos only had the *EYS^wt^* allele and flanking maternal SNPs (**Table S1**), and 2 had indels on the paternal allele. At the blastocyst stage (n=3 embryos), 2 showed only the *EYS^wt^* allele (**Table S1**), which was confirmed by on-target NGS and allelic discrimination qPCR, and one showed an indel (**Table S2, Table S4**). Therefore, of 7 embryos analyzed with Sanger genotyping and on-target NGS, 4 (57%) were *EYS^wt^* embryos (**Fig. 3B**).

**Figure 3:**
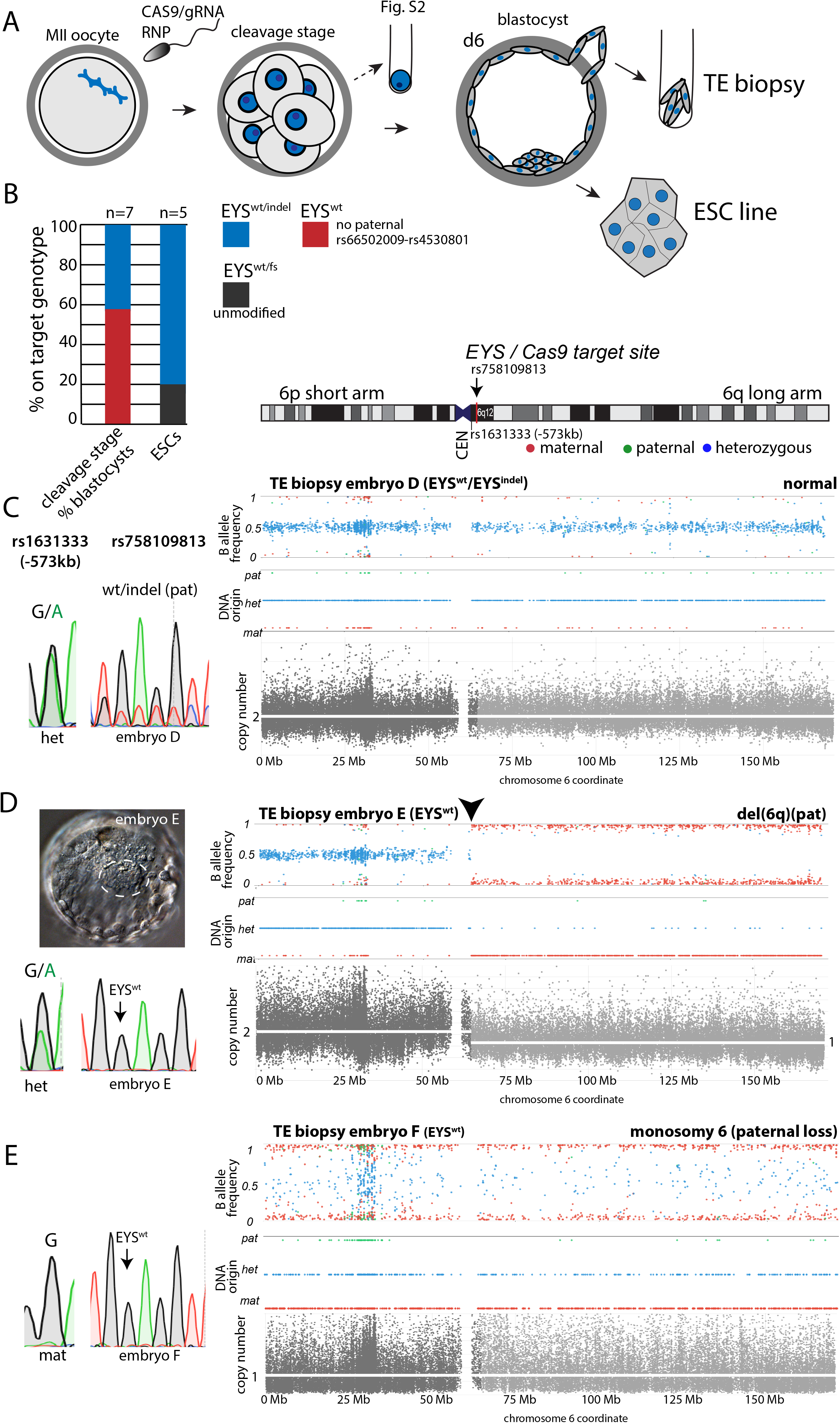
Chromosome loss in embryos with a ‘wild-type’ genotype. **A**) Schematic of ICSI at the MII stage with Cas9 RNP followed by development to the cleavage and blastocyst stages. Analysis at the cleavage stage involves harvesting of all cells and is incompatible with further development, while trophectoderm biopsy at the blastocyst stage is compatible with ESC derivation. **B)** On-target analysis by Sanger sequencing for embryos and embryonic stem cells. Shown is the percentage of embryos with the indicated genotypes. **C-E)** Heterozygosity analysis on chromosome 6 by SNP array for indicated samples, and Sanger sequencing profiles for the mutation site and a SNPs informative of parental origin centromeric to the cut site in the same embryo. On top, shown is the chromosomal location of the Cas9 target site at the EYS locus. The SNP array plots show allele frequency (plot 1) relative to a reference B allele, and parental origin of the detected SNP (plot 2). Only SNPs in which the maternal genotype was homozygous for one allele (red) and the paternal genotype was homozygous for the other allele (green) were used for analysis. Blue indicates a heterozygous (normal) embryo genotype. Plot 3 (grey) indicates copy number through signal intensity quantification, with flanking sides of rs758109813 shaded in lighter or darker grey. **C**) Blastocyst with a heterozygous indel which also gave rise to an ESC line. **D)** Blastocyst shown with its *EYS^wt^* Sanger genotype, as well as blastocyst morphology. Dotted line outlines the inner cell mass (ICM). **E**) Analysis of a trophectoderm of another blastocyst. No ESC lines were obtained from the ICM.

Embryos with only maternal sequences at the mutation site could arise by interhomolog recombination giving rise to *EYS^wt/wt^*, which may also lead to loss of linked paternal SNPs. Alternatively, paternal sequences could simply be lost (*EYS^wt^*) due to inadequate repair of the Cas9-induced DSB. To determine whether an *EYS^wt/wt^* genotype could be obtained, 10 blastocyst stage embryos, 3 of which had been genotyped by Sanger sequencing, were used for ESC derivation and 5 ESC lines were obtained. Four showed an indel at the paternal mutation site, while one maintained an unmodified paternal allele (**Fig. 3B**. **Table S1**). The derived stem cell lines were karyotypically normal (**Fig. S1, Table S3**), and showed no evidence of off-target activity at three top off-target sites. Despite the presence of a distinct inner cell mass, no ESC lines were obtained from two *EYS^wt^* blastocysts, which only had the wild-type maternal allele by TE biopsy. Instead, both plated blastocysts underwent cell death during attempted derivation, suggesting that these cells contained a lethal abnormality. Thus, there was no evidence for interhomolog repair after MII injection.

### Segmental and complete loss of the paternal chromosome

The loss of the paternal allele, including at SNPs distant from the mutation site, together with a lack of their representation in ESC lines with only the *EYS^wt^* allele, led us to investigate whether large deletions or chromosomal rearrangements accounted for the loss. For this, we first performed genome-wide SNP array analysis of genomic DNA from the sperm and oocyte donors to identify donor-specific alleles. Only identifying SNPs which were homozygous within the donors but different between the donors were subsequently used for analysis of embryos and stem cell lines.

Analysis of ESC lines showed heterozygosity for parent-of-origin-specific SNPs along the entire chromosome 6 and throughout the genome due to the presence of maternal and paternal chromosomes (**Fig. S1, Table S3**). We next controlled for the effect of genome amplification and analyzed trophectoderm biopsies and single human ESCs, since amplification of genomic DNA can alter the normal allelic ratio from 1:1 due to the stochasticity of amplification, in particular when only one cell is used. In both single amplified ESCs and trophectoderm biopsies of embryos that gave rise to the stem cell lines (**Fig. S1A**), heterozygous alleles were identified along the entire chromosome 6 and throughout the genome, albeit at a wider distribution (compare **Fig. S1A** and **Fig. S1C** with **Fig. S1B**). For some SNPs, only one of the alleles reaches significance of detection and thus, both paternal alleles (green dots), maternal alleles (red dots), as well as heterozygous alleles (blue dots) are called throughout chromosome 6, confirming the presence of both chromosomes. As a control, we also analyzed amplified genomic DNA from cumulus cells of an egg donor, which showed homozygous maternal alleles along chromosome 6 and throughout the genome (**Fig. S1D**). These controls provide a reference for the expected appearance of amplified genomic DNA in embryo samples, and the presence or absence of paternal chromosome 6.

We then examined TE biopsies from blastocyst embryos that had been genotyped at *EYS* by Sanger sequencing. Embryo D, which carried a heterozygous indel, showed heterozygosity and uniform signal intensity across chromosome 6 (**Fig. 3C**). In contrast, one of the two *EYS^wt^* blastocysts which had failed to develop a stem cell line (embryo E) showed loss of heterozygosity spanning the region from the Cas9 cut site at *EYS* all the way to the telomere of the long arm of paternal chromosome 6 (**Fig. 3D**). The loss of heterozygosity at the *EYS* break site was paralleled by a loss of signal intensity due to copy number loss along the entire length of the chromosome arm 6q (**Fig. 3D**). Therefore, loss of genetic material, rather than break-induced replication from the paternal centromere to the telomere is responsible for the absence of paternal SNPs. Furthermore, the other *EYS^wt^* blastocyst (embryo F) showed monosomy for the maternal chromosome 6, indicating loss of the entire paternal chromosome 6 (**Fig. 3E**). Both blastocysts were euploid for other autosomes (**Table S3**).

To better understand the process of chromosome loss, we performed analysis of chromosome content in blastomeres at the cleavage stage. We harvested 23 individual blastomeres from 4 embryos (**Table S1**). Blastomeres (n=11) from two cleavage stage embryos had a maternal-only genotype at *EYS*, which was confirmed by on-target NGS (**Table S2**). All four blastomeres analyzed from one embryo using SNP arrays showed segmental rearrangement of paternal chromosome 6. Of the 4 cells, 3 had losses of chromosome 6q distal of the Cas9 cleavage site, as seen by SNP array, copy number analysis and by Sanger genotyping of parent-of-origin-specific SNPs 10kb distal to the break site (**Fig. S2A**, **Table S1**). In one cell, chromosome 6q was gained, resulting in an overrepresentation of paternal SNPs distal of *EYS*, and an increase in copy number. Therefore, a chromosome 6q segment that was lost in one blastomere missegregated to the other in mitosis. Despite the gain of paternal chromosomal material, the on-target genotyping analysis was *EYS^wt^*, suggesting that these are paternal chromosome segments with an unrepaired break at *EYS*.

The other embryo had the *EYS^wt^* allele at the mutation site in 5/8 blastomeres, two blastomeres had no on target genotype, and one had an indel on an *EYS^wt^* allele (**Fig. S2B**, **Table S1**). Again, we found complementary losses and gains of paternal chromosome 6q. Four cells were monosomic for the maternal chromosome, one cell contained a gain of paternal chromosome 6q, and two cells contained only paternal chromosome 6p and were chaotic aneuploid. Therefore, Cas9 cleavage resulted in different chromosome 6 content even in cells with the same on-target genotype *EYS^wt^*.

Blastomeres from the remaining two cleavage stage embryos showed repair by end joining of the paternal allele. One embryo (embryo B) showed a uniform deletion of 5bp in all (4/4) blastomeres. In the other embryo, 4/5 blastomeres (embryo A) showed a net 2bp indel and one cell showed a wild-type only genotype by Sanger sequencing, as well as by on-target NGS (**Table S1**). Blastomeres from all *EYS^wt/indel^* heterozygous cells showed heterozygosity and balanced copy number on chromosome 6 (**Fig. S3**). One *EYS^wt^* blastomere had monosomy 6 due to paternal loss, and another had a segmental deletion at chromosome 6q21 on the maternal chromosome, 40Mb telomeric of the *EYS* locus. Though these cleavage stage embryos had numerous mitotic aneuploidies on other autosomes (**Table S3**), these may be spontaneous and representative of the frequent mitotic abnormalities present in cleavage stage embryos (Vanneste et al., 2009).

Of the 14 biopsies (12 blastomeres and 2 trophectoderm biopsies) with *EYS^wt^* genotypes from 5 different embryos, all showed segmental or whole chromosome aneuploidies of paternal chromosome 6 (**Table S1**). Surprisingly, monosomy for either the long arm of chromosome 6 or the entire chromosome 6 was compatible with the development to the expanded blastocyst stage (**Fig. 3D**). Thus, a common outcome of Cas9 RNP injections into MII oocytes is loss of the paternal allele due to segmental and whole chromosome loss, which appear as *EYS^wt^* cells in embryos characterized by on target sequencing (**Table S1**). These aneuploid cells were unable to produce pluripotent stem cell lines; only embryos with heterozygous indels gave rise to stem cells, even as these two genotypes each represented approximately half of preimplantation embryos (**Fig. 3B**). Recombination between homologous chromosomes indicated by novel combinations of maternal and paternal alleles within the same PCR product spanning rs6652009 to rs4530841 was not observed (**Table S1**).

### Cas9 RNP injection at the 2-cell stage results in chromosome loss after a single mitosis

An obstacle to repairing the paternal allele through recombination between homologous chromosomes at the zygote stage is that paternal and maternal genomes are packaged in two separate nuclei. To determine whether the maternal genome could provide a template for DSB repair when present with the paternal genome in the same nucleus, we injected Cas9 RNP into both cells of 13 two-cell stage embryos heterozygous for the *EYS* mutation and flanking SNPs (**Fig. 4A**). Single blastomeres were harvested and amplified from 9 cleavage stage embryos, and 4 embryos were analyzed at the morula and blastocyst stages on day 5 after fertilization using biopsies of multiple cells (**Fig. 4A**). Embryos dissociated to single blastomeres showed mosaicism for multiple alleles (**Fig. 4B):** of a total of 45 genotyped single cells and day 5 biopsies, 25 (55%) showed an *EYS^wt^* maternal-only genotype at the mutation site. Of these 25 samples, 22 were maternal only at the SNPs contained within the same PCR product, rs66502009 and rs12205397, and 13 were maternal only also at all the flanking SNPs 10kb proximal and distal of the Cas9 cleavage site, while 9 showed heterozygosity only at the 10kb distant flanking SNPs (**Table S1**). As Sanger sequencing is susceptible to allele-dropout, we performed on-target NGS on *EYS^wt^* blastomeres from 5 embryos and confirmed the *EYS^wt^* genotype (**Fig. 4C**, **Table S1**). In two *EYS^wt^* morulas harvested as biopsies of 5-15 cells, an edited paternal allele was present with reduced representation compared to the maternal allele due to mosaicism (**Fig. 4C**). Interestingly, of the 25 *EYS^wt^* samples, 3 cells from two different embryos showed flanking paternal SNPs on either side, which are captured within the same PCR product as *EYS^wt^* (**Fig. S4A, B**). NGS only detected a paternal allele at rs66502009, and only the maternal allele at rs758109813 (**Fig. 4D**, **Table S1**). SNPs of both maternal and paternal origin were detected throughout chromosome 6 by array analysis, demonstrating that this cell contained both paternal and maternal chromosome 6 (**Fig. S4C**). The novel linkages of maternal and paternal SNPs are possible interhomolog repair events, which occurred at a frequency of ~7% (3/45).

**Figure 4.**
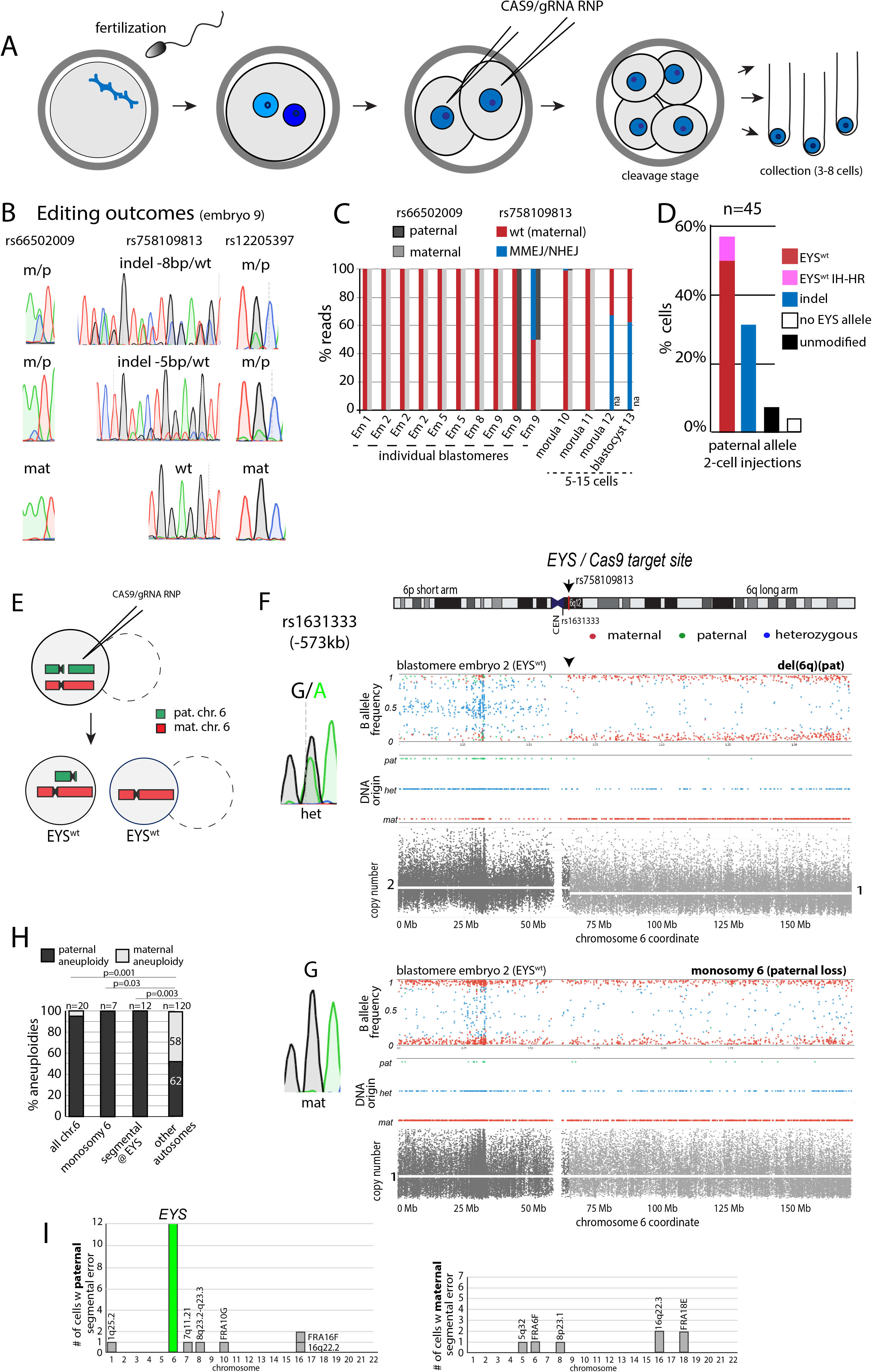
Chromosome loss and mosaicism after Cas9 RNP injection into two-cell stage embryos. **A)** Schematic of the experiment. A human oocyte is fertilized with EYS^2265fs^ mutant sperm, and is injected with Cas9 RNP only at the 2-cell stage, at 30-35h post ICSI, followed by analysis of individual cells at the cleavage stage. **B)** Sanger sequencing profiles of three different blastomeres (of 8 total) of the same embryo (embryo 9), showing mosaic outcomes. **C**) On-target NGS of embryo samples at the mutation site (rs758109813) and the linked SNP rs66502009, at the indicated stages. na=not applicable for samples without identifying parental SNP rs6652009. Em=embryo. **D**) Quantification of the percentage of cells with indicated genotypes determined by Sanger sequencing. No *EYS* allele represents samples without an *EYS* rs758109813 allele, but with genotypes of distant flanking SNPs. **E-G**) Analysis of sister blastomeres of an embryo after a single cell division post RNP injection. **E**) Schematic of the cell division products observed after a single cell cycle post Cas9 RNP injection. Two cells were successfully analyzed by SNP array, the cleavage products of the second injected cell that did not divide (dotted line), was not. (**F,G**). SNP arrays and Sanger sequencing of SNP rs1631333 for sister blastomeres, both *EYS^wt^* in on-targeting sequencing but with different chromosome 6 content. Shown is the chromosomal location of the Cas9 target site at the EYS locus and the SNP rs1631333 informative of parental origin. Only SNPs in which the maternal genotype was homozygous for one allele (red) and the paternal genotype was homozygous for the other allele (green) were used for analysis. Blue indicates a heterozygous (normal) embryo genotype. The plots show allele frequency (plot 1) relative to a reference B allele, parental origin of the detected SNP (plot 2), and copy number through signal intensity quantification (plot3, grey). The area telomeric of the *EYS* gene is shaded in a lighter grey. CEN= centromere. Pat=paternal. Mat=maternal. **H**) Quantification of aneuploidies according to parental origin. Statistical analysis was performed using Fisher’s exact test. **I**) Number of cells with segmental aneuploidies of maternal or paternal origin for each chromosome 1-22.

Furthermore, from the 45 samples, 12 (32%) showed end-joining events on the paternal allele, 3 cells showed no change, and 2 cells had no allele call even as flanking SNPs were detected (**Fig. 4D**, **Table S1)**.

In the remaining 3 cells, we observed heterozygous indels on an allele with a wild-type SNP rs758109813. One such edit had also been observed in the MII injections in ‘embryo C’. Therefore, Cas9 RNP cut the maternal allele with an efficiency of 4/84 (~5%), while in the same samples, editing of the paternal allele was 78/84 (93%). Thus, the presence of the suboptimal PAM site GAG allowed some cleavage of the maternal allele in the embryo, though with much lower efficiency than on the paternal allele.

To determine whether *EYS^wt^* blastomeres were caused by paternal chromosome loss, we tested blastomeres of two embryos after a single mitosis post-Cas9/RNP injection for heterozygosity using SNP arrays (**Fig. 4E-G**, **Fig. S5**). In one embryo, two sister blastomeres with an *EYS^wt^* genotype showed loss of chromosome 6q and its sister cell showed monosomy 6 due to loss of the paternal chromosome (**Fig. 4F,G**). For a remaining third cell of the same embryo that had not divided upon injection and with an *EYS^wt^* genotype, SNP array analysis was not successful.

In the other embryo, we found complementary loss of either paternal chromosomal arm 6q or 6p plus the centromere with breakpoints at the *EYS* locus (**Fig. S5A-D**). The long arm that was lost in one blastomere was gained in the other, as seen by an increased representation of paternal SNPs in one of the two sister blastomeres on chromosome 6q (**Fig. S5C**) and reciprocal copy number gain and loss on the q or the p arm (**Fig. S5B,C**). In addition, one ‘cell’ contained only chromosome 6p, no signal from chromosome 6q, and no other genomic DNA (**Fig. S5D, Table S3**). The exclusion of chromosomal arms in cytoplasmic fragments may be one mechanism of their elimination from the embryo. The reciprocal losses and gains of chromosomal arms were also observed by Sanger sequencing of rs1631333, located 573kb from the Cas9 cleavage site towards the centromere (**Fig. S5B-D, Table S1**). The product(s) of another cell of the injected 2-cell embryo could not be determined. Loss of heterozygosity across the centromere as in Fig. S5B is inconsistent with copy-neutral mitotic recombination, providing further support that the loss of paternal alleles occurred through the loss of genetic material rather than interhomolog recombination. Thus, Cas9 RNP injection at the 2-cell stage can result in the loss of the paternal chromosome through missegregation in mitosis due to an unrepaired DSB at the *EYS* locus.

Considering karyotypes of all embryos from injections at fertilization or at the 2-cell stage, all cells and samples (19/19) with loss of EYS^2265fs^ and of the flanking paternal alleles showed paternal specific abnormalities on chromosome 6. The one exception was a zygote that was analyzed prior to the first cell division, which showed an *EYS^wt^* genotype, and heterozygosity across chromosome 6 (**Fig. S2C**). Though chromosomal aneuploidies are common in human cleavage stage embryos (Vanneste et al., 2009), aneuploidies acquired after fertilization on autosomes 1-5 and 7-22 equally affected both paternal and maternal chromosomes (53% vs. 47%) (**Fig. 4H**, **Table S3**). All but one chromosomal abnormality occurred during embryonic mitosis (**Table S3**). In contrast, aneuploidies of chromosome 6 were significantly enriched on the paternal chromosome for both segmental errors as well as whole chromosome loss. Segmental errors of either paternal or maternal origin were found on other autosomes with break points mapping to several locations, including to common fragile sites, which were likely due to spontaneous chromosome breakage (**Fig. 4I**). In particular, one breakpoint found on the maternal chromosome 6q mapped to FRA6 F telomeric of *EYS* (**Fig. S3A**, **Table S3**). Thus, Cas9-induced cleavage in human embryos results in segmental and whole chromosome errors beyond what is observed in normal development.

## Discussion

The genome bestowed at fertilization determines much of our health as adults. While genetic mutations may be corrected after birth using somatic gene therapy, the efficacy of this approach is dependent on the number of cells that can be edited and disease-causing mutations will still be passed on to the next generation. In contrast, gene editing in the embryo will alter the genome of all cells, including the germline. An important requirement for clinical application is the ability to predict outcomes. Mosaicism prevents inferring the genotype of the fetus from the trophectoderm, and is thus incompatible with clinical use. Therefore, the application of CRISPR/Cas9 prior to the first round of DNA replication in the embryo is meaningful. We injected CRISPR/Cas9 at fertilization with sperm, and found that in most, but not all instances, the outcome is non-mosaic editing. In 11/14 (79%) events detected at the zygote stage and in developing embryos (consisting of three uniform edits in androgenotes, five in isolated zygote nuclei, one in a 2PN zygote, one at the cleavage stage, and one in a TE biopsy and matching ESC line from the same blastocyst), a single modified paternal allele was present, and hence occurred before replication of *EYS*. However, as mosaicism was seen in 3/14 (21%) of embryos (two in androgenotes and one cell in a cleavage stage embryo), understanding the timing of DNA replication, as well as temporally controlled activity of Cas9 will likely be required for eventual clinical use.

Double-strand break repair occurred predominantly through nonhomologous end joining (NHEJ) or microhomology-mediated end joining (MMEJ). The placement of the guide RNA at *EYS^2265fs^* results in a cut between 3 bp of microhomology, leading to the elimination of 5 nucleotides through MMEJ as the most common repair event. The deletion of 5 bp restores the reading frame of the *EYS^2265fs^* allele through the loss of 2 amino acids relative to the wild-type allele in the first of five lamin AG domains (Abd El-Aziz et al., 2008). The reading frame was also restored through the insertion of a single A nucleotide by NHEJ, resulting in two amino acid substitutions relative to the wild-type allele. These recurrent 1bp insertions are the consequence of filling in 1bp overhangs created by Cas9 cutting (Jasin, 2018; Lemos et al., 2018). As disease causing missense mutations in *EYS* map to the 4^th^ and 5^th^ lamin AG domains (Khan et al., 2010), restoring the frame may also restore function. Additional functional testing of the altered protein in an animal model, such as zebrafish or flies, will be required prior to considering clinical use in gene therapy.

Surprisingly, a frequent outcome of a single Cas9-induced double-strand break in human embryos at the *EYS* locus is the loss of part or all of paternal chromosome 6. We found loss of the long arm 6q, the short arm 6p, as well as of the entire chromosome in blastomeres and trophectoderm biopsies. Embryos with an *EYS^wt^* genotype that were tested for aneuploidies using SNP arrays showed aneuploidies of chromosome 6; by contrast, embryos with heterozygous edits *EYS^wt/indel^* did not. While segmental aneuploidies with a breakpoint at the *EYS* locus are readily attributed to Cas9 cleavage, whole chromosome errors can occur by either mitotic errors or when uniform in all cells of the embryo, through meiotic segregation errors. However, meiotic chromosome losses are predominantly of maternal origin, and spontaneous losses of chromosome 6 are relatively infrequent (Franasiak et al., 2014). In contrast, in our samples, whole chromosome losses were of paternal origin, and occurred after fertilization. Different cells of the same embryo showed both whole chromosome loss, as well as mirroring losses of the long or the short arms of chromosome 6. Therefore, pericentromeric cleavage at the *EYS* locus destabilizes the entire chromosome and results in both segmental as well as whole chromosome loss.

Prior to mitosis, an unrepaired break leads to an undetectable paternal allele, which appears as *EYS^wt^* in an on-target sequencing assay due to the presence of the maternal allele, and euploid in an array or copy number analysis. In mitosis, the unrepaired break is converted to a segmental or whole chromosome loss, which again appear as *EYS^wt^* in an on-target sequencing result. This is consistent with the ability of human zygotes to enter mitosis with unresolved DNA damage (Chia et al., 2017) and suggests that checkpoint activation by a single DSB is insufficient to mediate cell cycle arrest at this stage of human development. The frequent entry of the human embryos to repair a DSB prior to mitosis, raises the intriguing possibility that failure to repair endogenous DSBs may contribute to the frequent mosaic aneuploidies seen in human embryos (Kort et al., 2015). Though the location of segmental aneuploidies on other autosomes do not co-localize with top off-target sites, we cannot currently exclude that Cas9 cleavage elsewhere in the genome may also have caused some of the aneuploidies observed in cleavage stage embryos. Despite the ultimately lethal loss of chromosome 6, blastocysts with normal morphology can form, but fail to establish a stem cell line. Indeed, preimplantation embryos are tolerant of monosomies for development until the blastocyst stage (Franasiak et al., 2014), but do not provide monosomic ES cell lines (Lavon et al., 2008). The loss of chromosome 6 may be compensated by maternal products provided in the egg until ~ day 6 of development.

Our results serve as a cautionary note for the use of induced double-strand breaks in editing the genome of human embryos for clinical use. Chromosomal material may be lost and result in developmental abnormalities due to aneuploidy. Our findings are likely relevant to the interpretation of an earlier study using a guide RNA targeted to the MYBPC3 locus located on chromosome p11.2 near the centromere (Ma et al., 2017). Unrepaired breaks and the loss of the chromosomal arm or the entire chromosome may have contributed to the loss of the mutant paternal allele, resulting in the detection of only a wild-type maternal allele by on-target sequencing. The karyotype of these embryos was not determined. Another study found no evidence for whole chromosome aneuploidies after Cas9 cleavage at POU5F1, a locus on the short arm of chromosome 6 at p21.3 (Fogarty et al., 2017). It will be interesting to determine the incidence of segmental aneuploidies in these embryos and whether the outcomes of a DSB - segmental or whole chromosome loss - depend on chromosomal location is not currently known. While *POU5F1* is located on the short arm of chromosome 6 at p21.3, *EYS* is pericentromeric, which may have different consequences for centromere function and induction of whole chromosome or segmental loss.

Our study suggests that a DSB induced by even a single gRNA can result in the allele-specific removal of a chromosome in human embryos. Previous studies in mouse embryos demonstrated elimination of sex chromosomes by targeting Cas9 to centromeric repeats of the Y chromosome, or to multiple locations on the same chromosome (Adikusuma et al., 2017; Zuo et al., 2017). The finding that a single Cas9 induced cut can result in such outcome in human embryos suggests species-specific differences in repair or checkpoint control. Such induction of allele-specific chromosome loss may have clinical applications. Aneuploidies caused by abnormal meiosis are common in human oocytes, which is a major obstacle to fertility treatments and a cause for congenital abnormalities (Hassold and Hunt, 2001). Monosomic chromosomal gains are estimated to occur in about 5% of human oocytes (McCoy et al., 2015), which may be amenable to correction by Cas9. The placement of one or more gRNAs on both sides of the centromere may further increase the frequency of whole chromosome loss and thereby allow efficient allele-specific correction of trisomies in human embryos. Prior to considering clinical use, improvements in the specificity and timing of Cas9 cleavage will be required, as well as a better understanding how chromosome fragments are lost from the developing embryo, and whether fragments can integrate elsewhere in the genome. In a basic research context, the results presented here show that Cas9 provides a powerful tool to understand DNA repair in the human preimplantation embryo and the genetic and developmental consequences of DSBs.

## Methods

### Gamete donation

Oocyte donors were recruited from subjects participating in the Columbia Fertility oocyte donor program and were provided the option to donate for research instead of for reproductive purpose. Oocyte donors underwent controlled ovarian stimulation and oocyte retrieval as in routine clinical practice and as previously described (Zakarin Safier et al., 2018). Oocyte donors also provided a vial of blood (3-5ml) for isolation of genomic DNA and a skin biopsy. 67 Oocytes from a total of 8 different oocyte donors were used. Age at donation was 27-31. Oocytes were cryopreserved using Cryotech vitrification kit until genotyping of the oocyte donor at mutation site and flanking SNPs was completed and then thawed using the Cryotech warming kit. The sperm donor provided material via Male-FactorPak collection kit (Apex Medical Technologies MFP-130), which was cryopreserved using washing medium from MidAtlantic Diagnostics (ART-1005) and TYB freezing media from Irvine Scientific (90128). All gamete donors provided signed informed consent. All human subjects research was reviewed and approved by the Columbia University Embryonic Stem Cell Committee and the Institutional Review Board. All human embryos were cultured for 1-6 days, in accordance with internationally accepted standards to limit developmental progression to less than 14 days (ISSCR, 2016).

### RNP preparation

Guide RNA 5’-G<u>UGUGUCUUUCUUCUGUACUG</u>GUUUUAGAGCUAGAAAUAGCAAGUUAAAAUAAG GCUAGUCCGUUAUCAACUUGAAAAAGUGGCACCGAGUCGGUGCUUUU-3’ was obtained from Synthego, with the target RNA underlined. EnGen Cas9 NLS, S. pyogenes was obtained from NEB (M0646T). For ribonucleoprotein (RNP) preparation 3.125 μL of 20 μM Cas9 and 0.776 μL of 100 μM sgRNA was combined and incubated at room temperature for 10 minutes, followed by addition of 46μl injection buffer consisting of 5mM Tris-HCl, 0.1mM EDTA, pH 7.8.

### Oocyte manipulations

All manipulations were performed in an inverted Olympus IX71 microscope using Narishige micromanipulators on a stage heated to 37°C using Global Total w. HEPES (LifeGlobal LGTH-050). Oocyte enucleation for androgenesis was performed as previously described (Sagi et al., 2019; Yamada et al., 2014). Briefly, oocytes were enucleated in 5μg/ml cytochalasinB (Sigma-Aldrich C2743), the zona pellucida was opened using a zona laser (Hamilton Thorne) set at 100% for 300μs. The spindle was visualized using microtubule birefringence and removed using a 20μm inner diameter Piezo micropipette (Humagen). Intracytoplasmic sperm injection was identical for both nucleated and enucleated metaphaseII oocytes. Sperm was thawed to room temperature for 10 minutes and transferred to a 15mL conical tube. Quinn’s Sperm Washing Medium (Origio) was added dropwise to a final volume of 6mL. The tube was then centrifuged at 300x g for 15 minutes. Supernatant was removed and an additional wash was performed. Upon removal of supernatant from second wash, pellet was suspended in wash media and analyzed for viability.

Manipulation dishes consisted of a droplet with 10%PVP, a 10-20μl droplet with RNP in injection buffer, and a droplet of Global Total w. HEPES. Sperm was mixed with 10%PVP (Vitrolife), and individual sperm was immobilized by pressing the sperm tail with the ICSI micropipette (Humagen), picked up and transitioned through the RNP droplet before injection. After all manipulations, cells were cultures in Global total (LifeGlobal) in an incubator at 37°C and 5% CO2. Pronucleus formation was confirmed on day 1 after ICSI. 2-cell injections were performed at least 3 hours after cleavage, between 30-35h post ICSI. Earlier injection resulted in lysis. The tip of an injection needle was nicked and small amounts of the Cas9RNP was injected manually using a Narishige micromanipulator.

### Genome amplification and Genotyping

Zygotes were collected at 20h post ICSI, and single blastomeres on day3 to day4.Trophectoderm biopsies were obtained on day6 of development using 300ms laser pulses to separate trophectoderm from the inner cell mass. Single zygote nuclei were extracted from zygotes in the presence of 10mg/ml CytochalasinB and 1mg/ml nocodazole at 20h post ICSI. All samples were placed in single tubes with 2μl PBS. Amplification was performed using either Illustra GenomePhi V2 DNA amplification kit (GE Healthcare 95042-266), or REPLI-g single cell kit (Qiagen Cat #150345) according to manufacturer’s instructions. Genotyping was performed using primers for amplification and sequencing as listed in **Table S5**. PCR was performed using AmpliTaq Gold (ThermoFisher Cat. #4398886). TOPO-TA cloning (Thermo Fisher Cat #450640) was done for heterozygous edits and 5-10 colonies sequenced to identify the two edits individually (e.g. Fig. 1F).

For on-target NGS, primers were designed to amplify the specific region of the EYS gene. Once amplified, the PCR product was purified via sodium acetate precipitation. After purification samples were sent out for AmpliconEZ sequencing (Genewiz). Results were given in the format of raw reads and fastq files and an analysis of read frequency was conducted. Each sample yielded > 50,000 reads. Sequences with a read count that exceeded 0.2% of the total reads (>100 reads) were considered significant. Those that had less than 100 reads were considered insignificant due to the possibility that these were due to PCR artifacts or sequencing errors. Base changes were analyzed at the region of Cas9 cutting, and paternal and maternal alleles were called based on rs66502009, and rs758109813.

### EYS mutation allelic discrimination qPCR

A TaqMan assay (C_397916532_10) was used to obtain allelic discrimination results for the EYS mutation (rs758109813; NG_023443.2:g.1713111del) using a QuantStudio 3 instrument and following the manufacturer’s recommendations (ThermoFisher). See **Table S4** for results.

### Genome-wide SNP array

Embryo biopsies were amplified at Columbia University using either RepliG or GenomePhi as described above, or at Genomic Prediction Clinical Laboratory using ePGT amplification (Treff et al., 2019). Amplified DNA was processed according to the manufacturer’s recommendations for Axiom GeneTitan UKBB SNP arrays (ThermoFisher). Copy number and genotyping analysis was performed using gSUITE software (Treff et al., 2019). Parental origin of copy number changes was determined by genotype comparison of embryonic and parental SNPs. SNP array data are available at GEO under accession number <u>GSE148488</u>.

For copy number analysis, raw intensities from Affymetrix Axiom array are first processed according to the method described (Mayrhofer et al., 2016). After normalizing with a panel of normal males, the copy number is then calculated for each probeset. Normalized intensity is displayed. Mapping of endogenous fragile sites was done through visual evaluation of loss of heterozygosity.

### Derivation and culture of pluripotent stem cells

Stem cells were derived after trophectoderm biopsy and plating for the inner cell mass as previously described (Yamada et al., 2014). Two stem cell lines with the parental genotypes wt/EYS^2265fs^, eysES6 and eysES1, were obtained and used for experiments to determine mechanisms of repair after Cas9 mediated cleavage of the eys mutant allele. Stem cells were cultured with StemFlex (Thermo Fisher A3349401) media on Geltrex (Thermo Fisher A1413302). Upon reaching 70% confluency, cultures were passaged at a ratio of 1:10, or cryopreserved in a solution of freezing media containing 40% FBS (Company) and 10% DMSO (Sigma). Passaging was performed by TrypLE dissociation to small clusters of cells, and plated in media containing Rock inhibitor Y-27632 (Selleckchem S1049) was added to media and removed within 24-48 hours. For later passage cells (> passage 10), Rock inhibitor was omitted. Cas9 off-target analysis was performed by identifying 3 top off-target sites using Cas-OFF finder online tool (http://www.rgenome.net/cas-offinder/), selecting SpCas9 of Streptococcus pyogenes. PCR primers for off-target analysis are indicated in **Table S5**.

### Double-strand break repair study in EYS^wt^/EYS^2265fs^ pluripotent stem cells

For CRISPR-Cas9 gene editing, cells were dissociated to single cells, and cells were nucleofected with Cas9-GFP (Addgene #44719), a gBlock (IDT) expression vector, using the Lonza Kit (VVPH-5012) and Amaxa Nucleofector. 1 million cells at 50% confluency at time of use were nucleofected per reaction using program A-023. GFP positive cells were sorted using BD SORP FACSAria cell sorters 48h post nucleofection at the Columbia Stem Cell Initiative Flow Cytometry Core. Cells were either directly harvested for genome amplification using identical methods as used for embryo blastomeres, and analyzed for gene editing, or plated onto Geltrex with StemFlex media with Rock inhibitor Y-27632 (Selleckchem). After 5-10 days, colonies grown from single cells were picked and harvested for DNA isolation and analysis. DNA collection from cultured cells was performed using QuickExtract (Lucigen QE09050).

## Supporting information

Zuccaro et al. supplemental

## Acknowledgments

We thank the New York Stem Cell Foundation and the Russell Berrie Foundation Program in Cellular Therapies for funding support, and Chyuan-Sheng Lin of the Columbia mouse transgenics core for advice with Cas9 RNP preparation and injection. We thank Daniela Georgieva and Rudolph L. Leibel for critical input on the manuscript and Qian Du for technical assistance. D.E. is a NYSCF - Robertson Investigator Alumnus.

## Author contribution

M.V.Z. and D.E. designed the study. M.V.Z. performed gamete preparations, C.M., E.W., R.K., D.E. and M.V.Z. performed PCR amplifications and Sanger sequencing. D.E. performed micromanipulation and biopsies. D.E. and M.V.Z. performed genome amplification and stem cell derivation. M.S. and M.V.Z. performed miseq analysis. C.M., E.W., and M.V.Z. performed CRISPR experiments in pluripotent stem cells. B.R., R.Z. and D.M. performed SNP array and qPCR. C.M. performed off-target site analysis. N.T. and J.X. analyzed SNP array and qPCR data. D.M. and N.T. performed karyotyping. R.L. recruited oocyte donors and retrieved oocytes. S.T. provided expertise on EYS and retinitis pigmentosa. M.J. contributed to experimental design and data interpretation. R.S.G. wrote human subjects research protocols and provided study information to the semen donor. D.E. and M.V.Z. wrote the paper with contributions from all authors.

**The authors declare no competing interests**

