## Supplementary material for "Reading frame restoration at the EYS locus, and allele-specific chromosome removal after Cas9 cleavage in human embryos": Zuccaro et al. supplemental

**Supplemental Material**

Content:

Supplemental Figures 1-5

Supplemental Tables 1-5

### Supplementary Figure 1

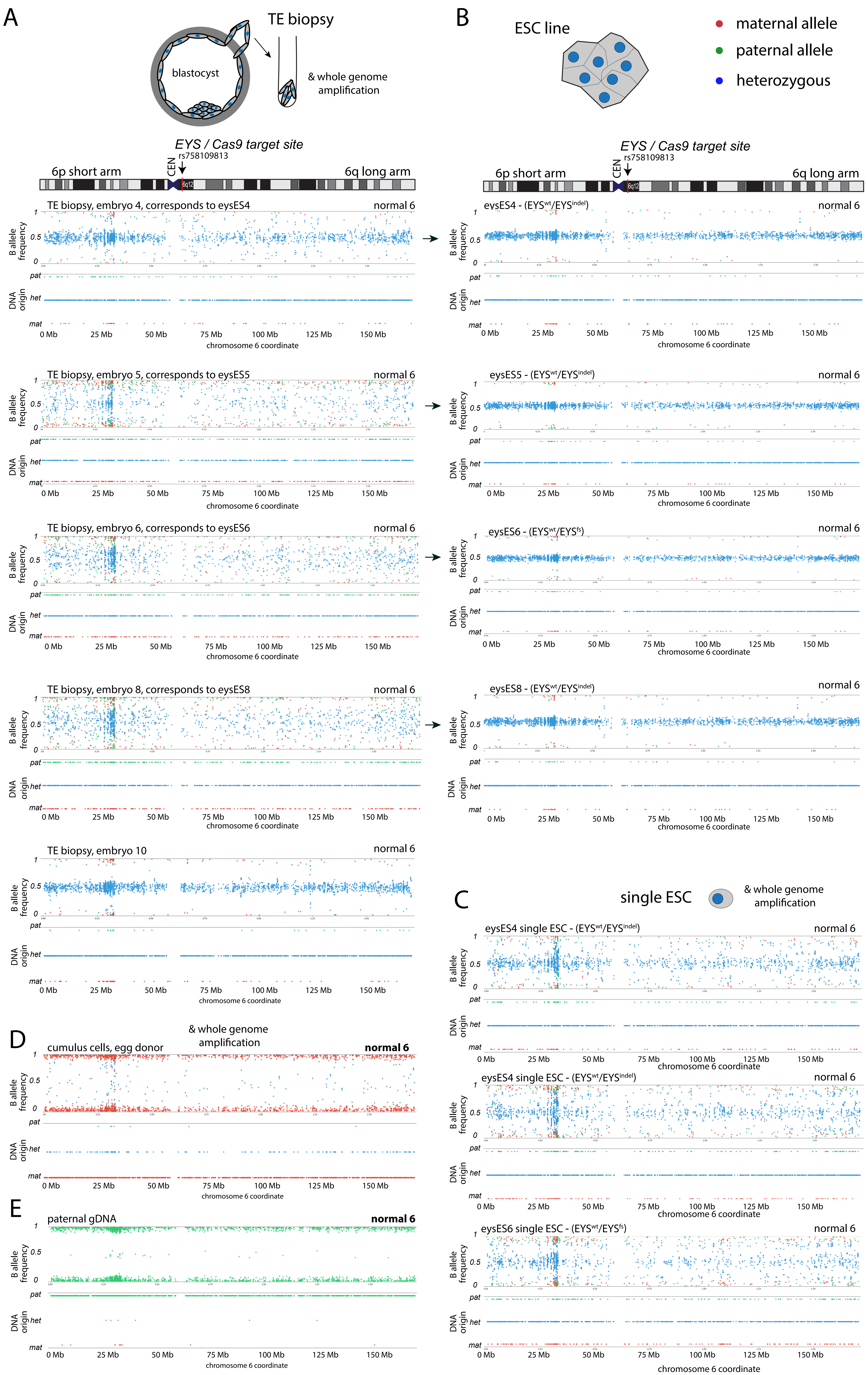

**Figure S1 | Heterozygosity in blastocysts and derived stem cell lines on chromosome 6 after Cas9 RNP injection at the MII stage**

For each panel, the developmental stage of the biopsy is indicated by a schematic at the top. The plots show allele frequency (plot 1) and parental origin of the detected SNP (plot 2). Only SNPs in which the maternal genotype was homozygous for one allele (red) and the paternal genotype was homozygous for the other allele (green) were included. Blue indicates a heterozygous (normal) embryo genotype. Arrow indicates the Cas9 cut site. **A**) Blastocyst biopsies. Each plot is a different blastocyst, four of which gave rise to a pluripotent stem cell line. The on-target genotype of the TE biopsies could not be determined. **B**) Chromosome 6 SNP array analysis of embryonic stem cell lines as a positive control for detection of heterozygosity along chromosome 6. **C**) The same stem cell lines are analyzed by SNP array after whole genome amplification from single cells. **D**) Analysis of amplified gDNA from cumulus cells of egg donor A as a control for a maternal-only chromosome 6 array profile. Increased signal intensity is seen in the HLA region of chromosome 6, which is likely an artifact. **E**) Analysis of paternal genomic DNA as a control for a paternal-only profile.

Related to Figure 3. For SNP array analysis of other autosomes see Table S3.

### Supplementary Figure 2

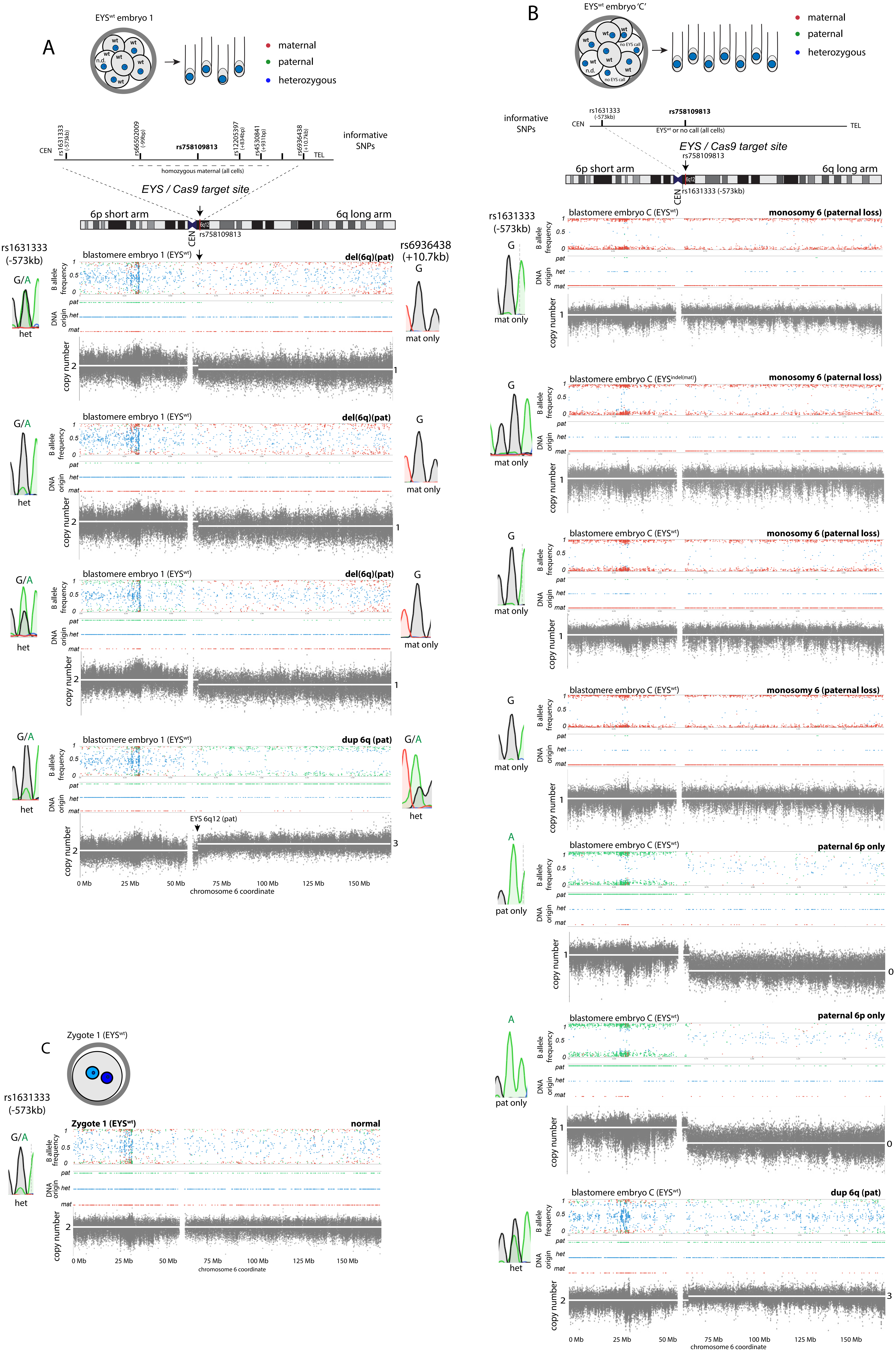

**Figure S2 | Chromosomal rearrangements at the EYS locus in blastomeres after Cas9 RNP injection at the MII stage**

For each panel, the biopsied embryo is indicated by a schematic at the top, and the number of cells successfully isolated and analyzed by SNP array is indicated with the number of tubes. The plots show allele frequency (top plot 1), parental origin of the detected SNP (plot 2), and copy number (grey plot). Only SNPs in which the maternal genotype was homozygous for one allele (red) and the paternal genotype was homozygous for the other allele (green) were included. Blue indicates a heterozygous (normal) embryo genotype. Arrow indicates the Cas9 cut site. **A** and **B** indicates two different embryos. **C**) Zygote (zygote 1) with an $EYS^{wt}$ genotype analyzed at the 1-cell stage prior to the first mitosis. CEN= centromere. For SNP array analysis of other autosomes see Table S3. Related to Figure 3.

### Supplementary Figure 3

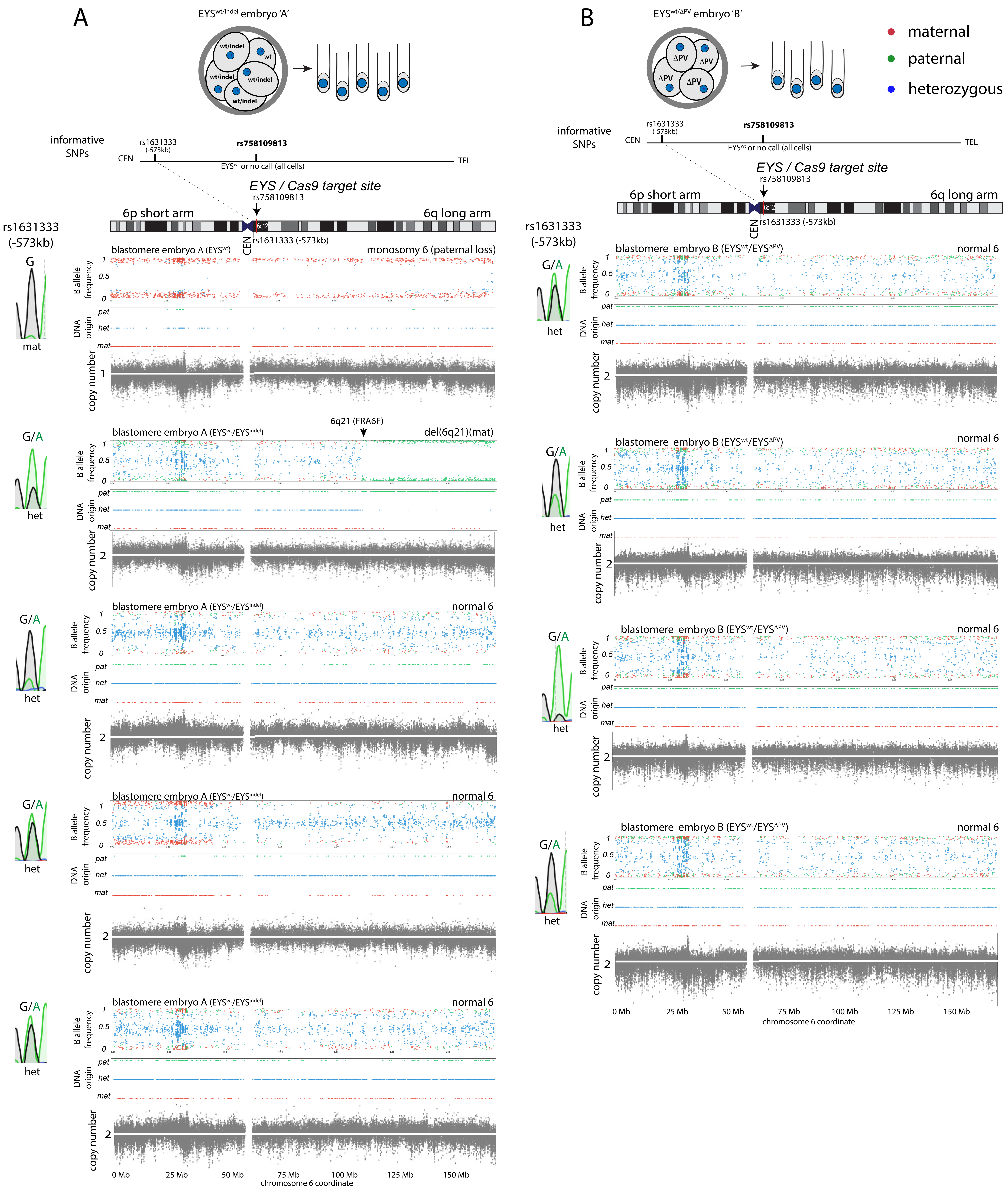

**Figure S3 | Heterozygosity in blastomeres with indels on chromosome 6 after Cas9 RNP injection at the MII stage**

For each panel, the biopsied embryo is indicated by a schematic at the top, and the number of cells successfully isolated and analyzed by SNP array is indicated with the number of tubes. The plots show allele frequency (top plot 1), parental origin of the detected SNP (plot 2), and copy number (grey plot). Only SNPs in which the maternal genotype was homozygous for one allele (red) and the paternal genotype was homozygous for the other allele (green) were included. Blue indicates a heterozygous (normal) embryo genotype. Arrow indicates the Cas9 cut site. **A** and **B** indicates each a different embryo. Representative copy number plots are shown. Note the segmental error of maternal origin on chromosome 6, at FRA6F. Different shades of grey indicate chromosomal regions distal and proximal to the EYS locus. CEN= centromere. For SNP array analysis of other autosomes see Table S3. Related to Figure 3.

### Supplementary Figure 4

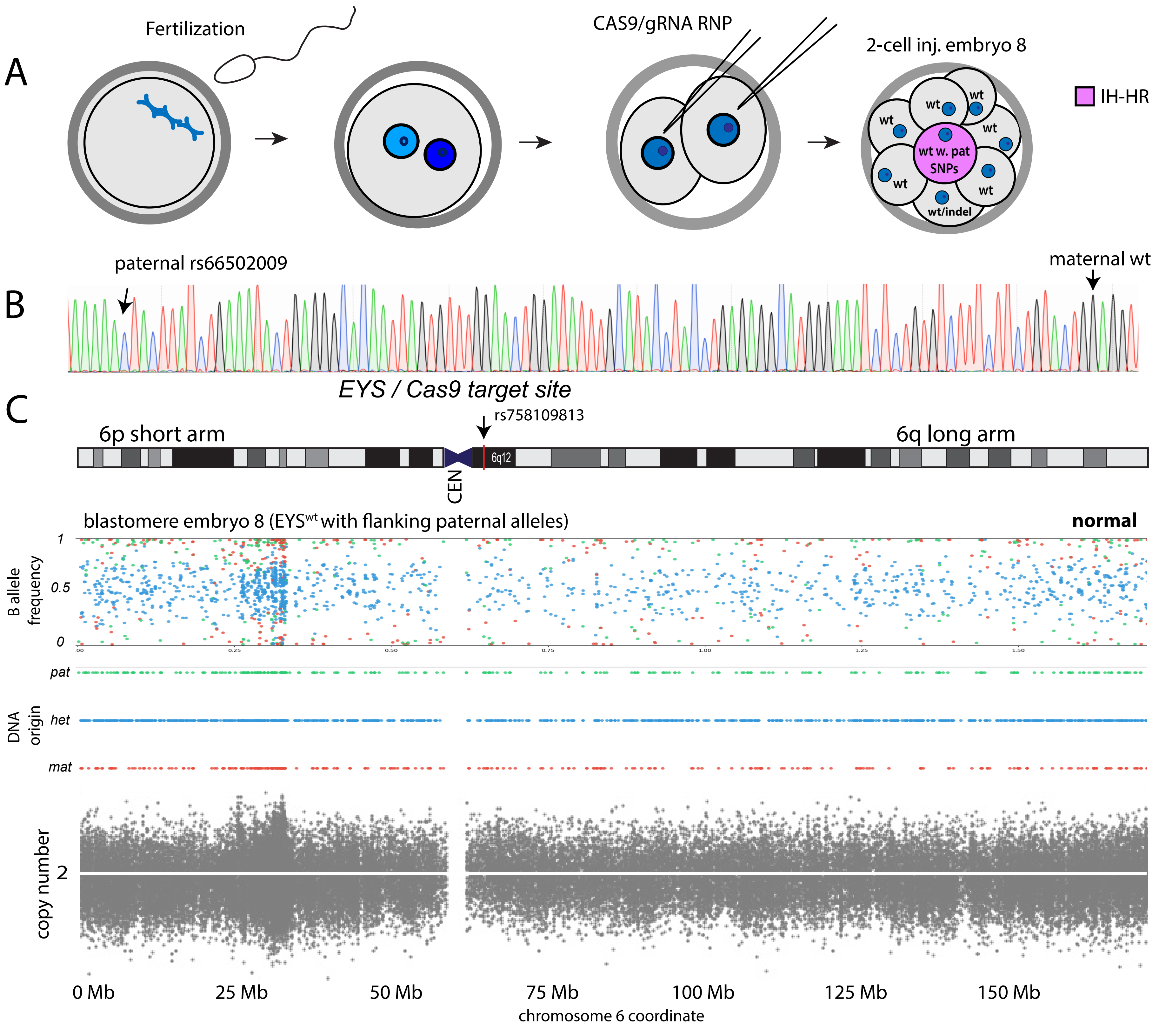

**Figure S4 | Possible interhomolog repair event after injection at the 2-cell stage**

**A)** Schematic of the experiment. Injection of Cas9/RNP at the 2-cell stage, one cell cycle after fertilization, and when paternal and maternal genomes are contained within the same nucleus.

**B**) Novel linkage of maternal $EYS^{wt}$ SNP and paternal rs66502009 shown by Sanger sequencing.

**C**) The SNP array plots shows allele frequency (top plot 1) and parental origin of the detected SNP (plot 2), and copy number analysis (plot 3, grey) for chromosome 6. Only SNPs in which the maternal genotype was homozygous for one allele (red) and the paternal genotype was homozygous for the other allele (green) were included.  Blue indicates a heterozygous (normal) embryo genotype. Arrow indicates the Cas9 cut site. The cell is heterozygous throughout chromosome 6, and carries an $EYS^{wt}$ only genotypes at the mutation site with paternal flanking SNPs. CEN= centromere. For SNP array analysis of other autosomes see Table S3. Related to Figure 4.

### Supplementary Figure 5

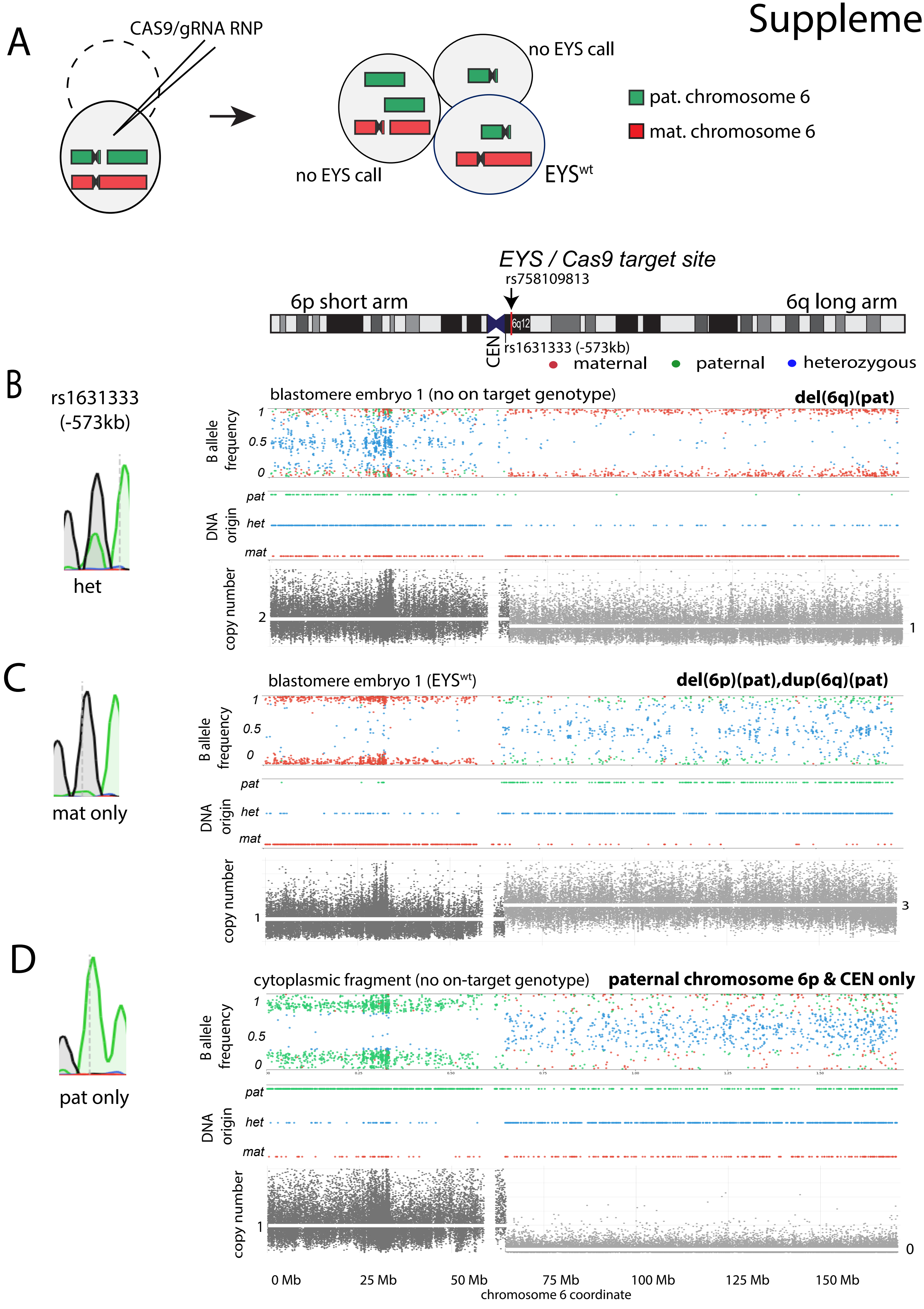

**Fig. S5 | Chromosome loss after Cas9 RNP injection into two-cell stage embryos**

**A)** Schematic of the cell division products observed after a single cell cycle post Cas9 RNP injection. Three cells were successfully analyzed, the cleavage products of the second injected cell was not (n.d., dotted line). (**B-D**). Sanger sequencing of rs1631333 and corresponding SNP arrays for 2 different cells (**B** and **C**) and one cytoplasmic fragment (**D**). Shown is the chromosomal location of the Cas9 target site at the EYS locus and the SNP rs1631333 informative of parental origin. The plots show allele frequency (plot 1) relative to a reference B allele, and parental origin of the detected SNP (plot 2). Only SNPs in which the maternal genotype was homozygous for one allele (red) and the paternal genotype was homozygous for the other allele (green) were used for analysis. Blue indicates a heterozygous (normal) embryo genotype. Plot 3 (grey) indicates copy number through signal intensity quantification. The area centromeric of the *EYS* gene and 6p are shaded in a dark grey, 6q telomeric of *EYS* in a lighter grey**.** **B)** Cell with a loss of chromosome 6q, **C**) cell with a loss of chromosome 6p and a gain of chromosome 6q, and **D**) 'cell' with only chromosome 6p without any other genomic DNA. Note that the cleavage products add up to 2 copies for each 6p and 6q arm.

CEN= centromere. Related to Fig. 4.

#### Supplemental Tables

**Table S1 | Integrated results from on target Sanger sequencing, on target deep sequencing, allelic discrimination qPCR, and karyotyping using SNP arrays.**

Light blue shaded areas are SNPs analyzed within a single PCR product. The flanking SNPs are each amplified by different primer sets. Alleles that are conclusively identifiable as of paternal origin are labeled in red. Homozygosity for a SNP is inferred through the presence of another heterozygous SNP within the same PCR product. SNPs in PCR products without heterozygosity, either due to homozygous alleles, or due to hemizygosity, which is indicated with a single letter. Nd= not determined. F= failed PCR amplification. NA= not applicable. Detailed results for on target sequencing are in Table S2, for karyotypes in Table S3, and for allelic discrimination PCR in Table S4.

**Table S2 | On target deep sequencing results at the *EYS* locus and flanking SNP rs66502009**

Detailed results from AmpliconEZ analysis.

**Table S3 | Results of SNP array analysis of embryonic cells after Cas9 cleavage and controls**

Detailed chromosomal content for all autosomes for indicated samples based on SNP array analysis. Paternal chromosome segregation errors are highlighted in green, maternal errors in orange.

**Table S4 | Allele discrimination quantitative PCR**

A TaqMan assay (C_397916532_10) was used to obtain allelic discrimination results for the EYS mutation (rs758109813; NG_023443.2:g.1713111del) using a QuantStudio 3 instrument and following the manufacturer's recommendations (Thermo Fisher). Shown are samples at the indicated stages.

**Table S5 | Oligonucleotides and primer sequences used for genotyping**

For the gBlock, the guide RNA sequence is colored in red.

### Table S1

| Cas9 RNP injection at | maternal egg genome origin | collection as | Chr6 (full kayotypes Table S4) | rs1631333 (-573kb) | rs1482455 (-11.2kb) | rs1482454 (-10.88kb) | rs9362339 (-10.71kb) | rs66502009 (-99bp) | rs66502009 (-99bp) deep sequencing (Table S3) | rs758109813 MUTATION SITE | rs758109813 MUTATION SITE deep sequencing (Table S3) | rs758109813 wt/fs allelic discrimination qPCR (Table S4) | rs12205397 (+834) | rs4530841 (+931) | rs34809101 (+10.34kb) | rs6936438 (+10.75kb) | altered allele sequence (@ rs758109813) | remarks |
|---|---|---|---|---|---|---|---|---|---|---|---|---|---|---|---|---|---|---|
| **Androgenesis @ 20h (no maternal genome)** | | | | | | | | | | | | | | | | | | |
| na | na | gDNA sperm donor | normal | na | T/T | G/G | T/T | C/C | C/C | fs/fs | fs/fs | fs/fs | A/A | A/A | -/- | A/A | GTCTTTCTTCTGTACTG-AGG | original paternal allele |
| MII androgenesis | na | 20h zygote | nd | nd | nd | nd | nd | C | nd | indel | nd | nd | A | A | nd | nd | CTT(-5bp)CTG-AGG | MMEJ -5bp |
| MII androgenesis | na | 20h zygote | nd | nd | nd | nd | nd | f | nd | f | nd | nd | f | f | f | f | na | failed amplification |
| MII androgenesis | na | 20h zygote | nd | nd | nd | nd | nd | C | nd | fs | nd | nd | f | f | nd | nd | na | unmodified |
| MII androgenesis | na | 20h zygote | nd | nd | nd | nd | nd | C | nd | indel/indel | nd | nd | A | A | nd | nd | CTGTAACTGAGG/CTT( -5bp)CTG-AGG | MMEJ -5bp & NHEJ RF restored |
| MII androgenesis | na | 20h zygote | nd | nd | nd | nd | nd | f | nd | f | nd | nd | f | f | f | f | na | failed amplification |
| MII androgenesis | na | 20h zygote | nd | nd | nd | nd | nd | C | nd | indel | nd | nd | A | A | nd | nd | GTA(-1bp)TG-AGG | NHEJ |
| MII androgenesis | na | 20h zygote | nd | nd | nd | nd | nd | C | nd | indel/indel | nd | nd | A | A | nd | nd | CTGTA(-1bp)TGAGG/CTT( -5bp)CTG-AGG | NHEJ/MMEJ -5bp |
| MII androgenesis | na | 20h zygote | nd | nd | nd | nd | nd | C | nd | indel | nd | nd | A | A | nd | nd | CTT(-5bp)CTG-AGG | MMEJ -5bp |
| **2PN individual pronuclei @ 20h** | | | | | | | | | | | | | | | | | | |
| na | na | gDNA sperm donor | normal | A/A | T/T | G/G | T/T | C/C | C/C | fs/fs | fs/fs | fs/fs | A/A | A/A | -/- | A/A | GTCTTTCTTCTGTACTG-AGG | original paternal allele |
| na | na | gDNA egg donor B | normal | G/G | C/T | A/G | C/T | A/C | nd | wt/wt | nd | wt/wt | G/A | G/A | nd | nd | GTCTTTCTTCTGTACTGGAGG | original maternal allele |
| MII - 1 | donor B | 20h zygote, individual nuclei | nd | G | T | G | T | C | nd | wt | nd | nd | nd | nd | - | A | na | |
| MII - 1 | na | 20h zygote, individual nuclei | nd | A | T | G | T | C | nd | indel | nd | nd | nd | nd | - | A | CTT(-5bp)CTG-AGG | MMEJ -5bp |
| MII - 2 | donor B | 20h zygote, individual nuclei | nd | G | T | G | T | f | nd | f | nd | nd | nd | nd | - | A | na | cut maternal allele |
| MII - 2 | donor B | 20h zygote, individual nuclei | nd | A | T | G | T | f | nd | indel | nd | nd | nd | nd | - | A | CTT(-5bp)CTG-AGG | MMEJ -5bp |
| MII - 3 | donor B | 20h zygote, individual nuclei | nd | G | T | G | T | C | nd | wt | nd | nd | nd | nd | - | A | na | na |
| MII - 3 | na | 20h zygote, individual nuclei | nd | A | T | G | T | C | nd | fs | nd | nd | nd | nd | - | A | na | unmodified |
| MII - 4 | donor B | 20h zygote, individual nuclei | nd | G | T | G | T | C | nd | wt | nd | nd | nd | nd | - | A | na | na |
| MII - 4 | na | 20h zygote, individual nuclei | nd | A | T | G | T | C | nd | indel | nd | nd | nd | nd | - | A | CTTTCTGCACACAGAGAGACTG-AGG | NHEJ |
| MII - 5 | donor B | 20h zygote, individual nuclei | nd | G | T | G | T | C | nd | wt | nd | nd | nd | nd | f | f | na | na |
| MII - 5 | na | 20h zygote, individual nuclei | nd | A | T | G | T | f | nd | f | nd | nd | nd | nd | - | A | cleaved paternal allele | no paternal on target genotype |
| MII - 6 | donor B | 20h zygote, individual nuclei | nd | G | T | G | T | C | nd | wt | nd | nd | nd | nd | - | A | na | na |
| MII - 6 | na | 20h zygote, individual nuclei | nd | A | T | G | T | nd | nd | indel | nd | nd | nd | nd | - | A | CTGTAACTG-AGG | NHEJ RF restored |
| MII - 7 | donor B | 20h zygote, individual nuclei | nd | G | nd | nd | nd | C | nd | wt | nd | nd | nd | nd | nd | nd | na | |
| MII - 7 | na | 20h zygote, individual nuclei | nd | A | T | G | T | C | nd | indel | nd | nd | nd | nd | nd | nd | CTG (-3bp) TG-AGG | |
| MII - 8 | donor B | 20h zygote, individual nuclei | nd | G | nd | nd | nd | C | nd | wt | nd | nd | nd | nd | nd | nd | na | |
| MII - 8 | na | 20h zygote, individual nuclei | nd | A | T | G | T | C | nd | f | nd | nd | nd | nd | nd | nd | cleaved paternal allele | |
| **control genotyping** | | | | | | | | | | | | | | | | | | |
| na | na | gDNA sperm donor | normal | A/A | T/T | G/G | T/T | C/C | C/C | fs/fs | fs/fs | fs/fs | A/A | A/A | -/- | A/A | GTCTTTCTTCTGTACTG-AGG | original paternal allele |
| na | donor G | gDNA egg donor G | nd | nd | C/C | A/A | C/C | A/A | A/A | wt/wt | wt/wt | nd | G/G | G/G | T/T | G/G | GTCTTTCTTCTGTACTGGAGG | original maternal allele |
| control - none | donor G | 2-cell d2 control embryo 1 | nd | G/A | C/T | A/G | C/T | A/C | nd | wt/fs | nd | nd | nd | nd | T/(-) | G/G | na | genotyping control |
| control - none | donor G | 2-cell d2 control embryo 1 | nd | nd | C/T | A/G | C/T | A/C | nd | wt/fs | nd | nd | G/A | G/A | T/(-) | G/G | na | genotyping control |
| na | donor A | gDNA egg donor A | normal | na | C/T | A/G | C/T | A/A | nd | wt/wt | nd | wt/wt | G/G | G/G | T/T | G/G | GTCTTTCTTCTGTACTGGAGG | original maternal allele |
| control - none | donor A | 2-cell d2 control embryo 2 | nd | nd | nd | nd | nd | A/C | nd | wt/fs | nd | nd | G/A | G/A | - | A | na | genotyping control |
| na | donor C | gDNA egg donor C | normal | nd | nd | nd | nd | A/A | nd | wt/wt | nd | wt/wt | G/G | G/G | nd | nd | GTCTTTCTTCTGTACTGGAGG | original maternal allele |
| control - none | donor C | single ESC eysES6 | normal | nd | nd | nd | nd | A/C | nd | wt/fs | nd | wt/fs | nd | nd | nd | nd | na | genotyping control |
| control - none | donor C | single ESC eysES6 | normal | nd | nd | nd | nd | A/C | nd | wt/fs | nd | nd | nd | nd | nd | nd | na | genotyping control |
| control - none | donor C | single ESC eysES6 | nd | nd | nd | nd | nd | A/C | nd | wt/fs | nd | nd | nd | nd | nd | nd | na | genotyping control |
| control - none | donor C | single ESC eysES6 | nd | nd | nd | nd | nd | A/C | nd | wt/fs | nd | nd | nd | nd | nd | nd | na | genotyping control |
| **2PN analysis @ 20h after MII injection** | | | | | | | | | | | | | | | | | | |
| na | na | gDNA sperm donor | normal | A/A | T/T | G/G | T/T | C/C | C/C | fs/fs | fs/fs | fs/fs | A/A | A/A | -/- | A/A | GTCTTTCTTCTGTACTG-AGG | original paternal allele |
| na | donor A | gDNA egg donor A | normal | G/G | C/T | A/G | C/T | A/A | nd | wt/wt | nd | wt/wt | G/G | G/G | T/T | G/G | GTCTTTCTTCTGTACTGGAGG | original maternal allele |
| MII | donor A | zygote 1 | normal | G/A | C | A | C | A | A (100%) | wt | wt | wt/not detected | G | G | T | G | cleaved paternal allele | loss of paternal allele |
| MII | donor A | zygote 2 | f | nd | T | G | T | A | A (100%) | wt | wt | wt/not detected | G | G | T | G | cleaved paternal allele | loss of paternal allele |
| na | donor B | gDNA egg donor B | normal | nd | C/T | A/G | C/T | A/C | nd | wt/wt | nd | wt/wt | G/A | G/A | nd | nd | GTCTTTCTTCTGTACTGGAGG | original maternal allele |
| MII | donor B | zygote 3 | nd | nd | C/T | A/G | C/T | nd | nd | wt/indel | nd | nd | G/A | nd | T/- | G/A | CTT(-5bp)CTG-AGG | MMEJ -5bp |
| **Blastomere collection after MII injection** | | | | | | | | | | | | | | | | | | |
| na | na | gDNA sperm donor | normal | A/A | T/T | G/G | T/T | C/C | C/C | fs/fs | fs/fs | fs/fs | A/A | A/A | -/- | A/A | GTCTTTCTTCTGTACTG-AGG | original paternal allele |
| na | donor A | gDNA egg donor A | normal | G/G | C/T | A/G | C/T | A/A | nd | wt/wt | nd | wt/wt | G/G | G/G | T/T | G/G | GTCTTTCTTCTGTACTGGAGG | original maternal allele |
| | | embryo 1 (7 cells) | | | | | | | | | | | | | | | | |
| MII | donor A | MII blastomere embryo 1 | f | f | f | f | f | A | A (100%) | wt | wt (100%) | f | G | G | f | f | na | loss of paternal allele |
| MII | donor A | MII blastomere embryo 1 | del 6q (pat) | G/A | T | G | T | A | A (100%) | wt | wt (100%) | wt/not detected | G | G | T | T | na | loss of paternal allele |
| MII | donor A | MII blastomere embryo 1 | del 6q (pat) | G/A | T | G | T | A | A (91%) / G (9%) | wt | wt (100%) | f | f | f | T | T | na | loss of paternal allele |
| MII | donor A | MII blastomere embryo 1 | dup 6q (pat) | G/A | T | G | T | A | A (100%) | wt | wt (100%) | wt/not detected | G | G | T/- | G/A | na | loss of paternal allele |
| MII | donor A | MII blastomere embryo 1 | del 6q (pat) | G/A | T | G | T | A | A (100%) | wt | wt (100%) | wt/not detected | G | G | T | T | na | loss of paternal allele |
| MII | donor A | MII blastomere embryo 1 | nd | G/A | T | G | T | A | nd | wt | nd | nd | G | G | T | T | na | loss of paternal allele |
| na | donor D | gDNA egg donor D | normal | G/G | T/T | G/G | T/T | C/C | nd | wt/wt | nd | wt/wt | A/A | A/A | -/- | A/A | GTCTTTCTTCTGTACTGGAGG | original maternal allele |
| | | embryo A (5 cells) | | | | | | | | | | | | | | | | |
| MII | donor D | MII blastomere embryo A | normal EYS locus, del6q21-27 (mat) | G/A | T | G | T | C/C | C (100%) | wt/indel | A(67%) / edited (33%) | wt/not detected | A | A | - | A | TTTCTTCT(-1bp)T(-1bp)CTG-AGG | NHEJ |
| MII | donor D | MII blastomere embryo A | normal | G/A | T | G | T | C/C | C (100%) | wt/indel | A(56%) / edited (44%) | wt/not detected | A | A | - | A | TTTCTTCT(-1bp)T(-1bp)CTG-AGG | NHEJ |
| MII | donor D | MII blastomere embryo A | normal | G/A | T | G | T | C/C | C (100%) | wt/indel | A(45%) / edited (55%) | wt/not detected | A | A | - | A | TTTCTTCT(-1bp)T(-1bp)CTG-AGG | NHEJ |
| MII | donor D | MII blastomere embryo A | normal | G/A | T | G | T | C/C | C (100%) | wt/indel | A(80%) / edited (20%) | wt/not detected | A | A | - | A | TTTCTTCT(-1bp)T(-1bp)CTG-AGG | NHEJ |
| MII | donor D | MII blastomere embryo A | monosomy 6 (pat) | G | T | G | T | C | C (100%) | wt | wt (100%) | wt/not detected | nd | nd | - | A | na | loss of paternal allele |
| | | Embryo B (4 cells) | | | | | | | | | | | | | | | | |
| MII | donor D | MII blastomere embryo B | normal | G/A | T | G | T | C/C | C (100%) | wt/indel | A(92%) / edited (8%) | wt/not detected | nd | nd | - | A | CTT(-5bp)CTG-AGG | MMEJ -5bp |
| MII | donor D | MII blastomere embryo B | normal | G/A | T | G | T | C/C | C (100%) | wt/indel | A(23%) / edited (77%) | wt/not detected | A | A | - | A | CTT(-5bp)CTG-AGG | MMEJ -5bp |
| MII | donor D | MII blastomere embryo B | normal | G/A | T | G | T | C/C | C (100%) | wt/indel | A(32%) / edited (68%) | wt/not detected | A | A | - | A | CTT(-5bp)CTG-AGG | MMEJ -5bp |
| MII | donor D | MII blastomere embryo B | normal | G/A | T | G | T | C/C | C (100%) | wt/indel | A(79%) / edited (21%) | wt/not detected | A | A | - | A | CTT(-5bp)CTG-AGG | MMEJ -5bp |
| | | Embryo C (8 cells) | | | | | | | | | | | | | | | | |
| MII | donor D | MII blastomere embryo C | nd | nd | nd | nd | nd | C | nd | wt | nd | nd | nd | nd | nd | nd | na | loss of paternal allele |
| MII | donor D | MII blastomere embryo C | monosomy 6 (pat) | G | nd | nd | nd | C | C (100%) | wt | wt (100%) | wt/not detected | nd | nd | nd | nd | na | loss of paternal allele |
| MII | donor D | MII blastomere embryo C | monosomy 6 (pat) | G | nd | nd | nd | C | C (100%) | wt | wt (100%) | wt/not detected | nd | nd | nd | nd | na | loss of paternal allele |
| MII | donor D | MII blastomere embryo C | paternal 6p only | A | f | f | f | f | na | f | na | no detection | nd | nd | f | f | no allele detected | loss of paternal allele |
| MII | donor D | MII blastomere embryo C | monosomy 6 (pat) | G | nd | nd | nd | C | C (100%) | wt | wt (100%) | wt/not detected | nd | nd | nd | nd | na | loss of paternal allele |
| MII | donor D | MII blastomere embryo C | paternal 6p only | A | f | f | f | f | na | f | na | no detection | f | f | f | f | no allele detected | loss of paternal allele |
| MII | donor D | MII blastomere embryo C | dup 6q (pat) | G/A | T | G | T | C | C (100%) | wt | wt (100%) | wt/not detected | A | A | nd | nd | na | loss of paternal allele |
| MII | donor D | MII blastomere embryo C | monosomy 6 (pat) | G | nd | nd | nd | C | nd | indel | nd | failed | nd | nd | nd | nd | CTTTCTTCT(-9bp )GTTTT | loss of paternal allele, altered maternal allele by MMEJ |
| **Trophectoderm Biopsy after MII injection** | | | | | | | | | | | | | | | | | | |
| na | na | gDNA sperm donor | normal (46,XY) | A/A | T/T | G/G | T/T | C/C | C/C | fs/fs | fs/fs | fs/fs | A/A | A/A | -/- | A/A | GTCTTTCTTCTGTACTG-AGG | original paternal allele |
| na | donor D | gDNA egg donor D | normal (46,XX) | G/G | T/T | G/G | T/T | C/C | C/C | wt/wt | nd | wt/wt | A/A | A/A | -/- | A/A | GTCTTTCTTCTGTACTGGAGG | original maternal allele |
| MII | donor D | TE biopsy d6 (embryo D) x2 | normal | G/A | T | G | T | C/C | nd | wt/indel | nd | wt/not detected | nd | nd | - | A | CTGTGTCTTACTG-AGG | NHEJ, two concordant biopsies |
| MII | donor D | TE biopsy d6 (embryo E) | del 6q (pat) | G/A | T | G | T | C | C (100%) | wt | wt (100%) | wt/not detected | nd | nd | - | A | na | loss of paternal allele |
| MII | donor D | TE biopsy d6 (embryo F) x2 | monosomy 6 (pat) | G | T | G | T | C | C (100%) | wt | wt (100%) | wt/not detected | nd | nd | - | A | na | loss of paternal allele, two concordant biopsies |
| *TE biopsies without Sanger genotype* | | | | | | | | | | | | | | | | | | |
| MII | donor C | TE biopsy (embryo 4) matches eysES4 x2 | normal | nd | nd | nd | nd | nd | nd | nd | nd | wt/not detected | nd | nd | nd | nd | nd | altered paternal allele, concordant with ICM |
| MII | donor C | TE biopsy (embryo 10) x2 | normal | nd | nd | nd | nd | nd | nd | nd | nd | wt/not detected | nd | nd | nd | nd | nd | altered paternal allele, defined paternal genotype not available |
| MII | donor C | TE biopsy( embryo 8) matches eysES8 | normal | nd | nd | nd | nd | nd | nd | nd | nd | wt/not detected | nd | nd | nd | nd | nd | altered paternal allele, concordant with ICM |
| MII | donor C | TE biopsy (embryo 5) matches eysES5 | normal | nd | nd | nd | nd | nd | nd | nd | nd | wt/not detected | nd | nd | nd | nd | nd | altered paternal allele, concordant with ICM |
| | | x2= two biopsies collected with consistent results | | | | | | | | | | | | | | | | |
| **ESC derivation after MII injection** | | | | | | | | | | | | | | | | | | |
| na | na | gDNA sperm donor | normal | A/A | T/T | G/G | T/T | C/C | C/C | fs/fs | fs/fs | fs/fs | A/A | A/A | -/- | A/A | GTCTTTCTTCTGTACTG-AGG | original paternal allele |
| na | donor C | gDNA egg donor C | normal | nd | nd | nd | nd | A/A | C/C | wt/wt | nd | wt/wt | G/G | G/G | nd | nd | GTCTTTCTTCTGTACTGGAGG | original maternal allele |
| MII | donor C | eysESC4 (from embryo 4) | normal | G/A | C/T | A/G | C/T | A/C | nd | wt/indel | nd | wt/not detected | G/A | G/A | T/- | G/A | CTGTTTTTCTTCTG-AGG | NHEJ |
| MII | donor C | eysESC5 (from embryo 5) | normal | nd | C/T | A/G | C/T | A/C | nd | wt/indel | nd | wt/not detected | G/A | G/A | T/- | G/A | CTGTCTTCTGTGTCTGTCTTT-8bpCTG-AGG | MMEJ plus 11bp insertion |
| MII | donor C | eysESC6 | normal | G/A | C/T | A/G | C/T | A/C | A/C | wt/fs | wt/fs | wt/fs | G/A | G/A | T/- | G/A | na | no change |
| MII | donor C | eysESC8 (from embryo 8) | normal | nd | C/T | A/G | C/T | A/C | nd | wt/indel | nd | wt/not detected | nd | nd | nd | nd | CTGAAACTG-AGG | NHEJ |
| na | donor D | gDNA egg donor D | normal | G/G | T/T | G/G | T/T | C/C | nd | wt/wt | nd | wt/wt | nd | nd | -/- | A | GTCTTTCTTCTGTACTGGAGG | original maternal allele |
| MII | donor D | eysESC9 | nd | G/A | nd | nd | nd | C/C | nd | wt/indel | nd | nd | nd | nd | nd | nd | CTGTGTCTTTCTG-AGG | NHEJ |
| na | donor I | gDNA egg donor I | nd | nd | C/C | A/A | C/C | A/A | nd | wt/wt | nd | nd | nd | nd | T/T | G/G | GTCTTTCTTCTGTACTGGAGG | original maternal allele |
| control - none | donor I | eysESC1 | nd | nd | C/T | A/G | C/T | A/C | nd | wt/fs | nd | nd | nd | nd | T/- | G/A | na | genotyping control |
| **2-cell stage injections** | | | | | | | | | | | | | | | | | | |
| na | na | gDNA sperm donor | normal | A/A | T/T | G/G | T/T | C/C | C/C | fs/fs | fs/fs | fs/fs | A/A | A/A | -/- | A/A | GTCTTTCTTCTGTACTG-AGG | original paternal allele |
| na | donor G | gDNA egg donor G | nd | nd | C/C | A/A | C/C | A/A | A/A | wt/wt | wt/wt | wt/wt | G/G | G/G | T/T | G/G | GTCTTTCTTCTGTACTGGAGG | original maternal allele |
| | | embryo 4 (4 cells) | | | | | | | | | | | | | | | | |
| 2-cell | donor G | embryo 4 | nd | nd | nd | nd | nd | A | nd | wt | nd | nd | G | G | T | G | na | loss of paternal allele |
| 2-cell | donor G | embryo 4 | nd | nd | C | A | C | A/A | nd | wt/ indel | nd | nd | G/G | G/G | T | G | CTTCTGTACCTGGAGG | loss of paternal allele, altered maternal allele NHEJ |
| 2-cell | donor G | embryo 4 | nd | nd | C/T | A/G | C/T | A/C | nd | wt/indel | nd | nd | G/A | G/A | T | G | GTA(-1bp)TG-AGG | NHEJ |
| na | donor A | gDNA egg donor A | normal | G/G | C/T | A/G | C/T | A/A | nd | wt/wt | nd | wt/wt | G/G | G/G | T/T | G/G | GTCTTTCTTCTGTACTGGAGG | original maternal allele |
| 2-cell | donor A | embryo 3 (1 cell) | nd | nd | f | f | f | A/C | nd | wt/fs | nd | nd | nd | nd | nd | nd | na | no change |
| | | embryo 7 (7 cells) | | | | | | | | | | | | | | | | |
| 2-cell | donor A | embryo 7 | nd | nd | C | A | C | C | nd | wt/indel | nd | nd | nd | nd | T | G | GTAACT(-1bp)AGG | NHEJ |
| 2-cell | donor A | embryo 7 | nd | nd | T | G | T | A/A | nd | wt/ indel | nd | nd | G/G | G/G | T | G | CTG(-1bp)ACTGGAGG | loss of paternal allele, altered maternal allele NHEJ |
| 2-cell | donor A | embryo 7 | nd | nd | T | G | T | A/C | nd | wt/indel | nd | nd | G/A | G/A | T | G | CTTC(-4bp)CTG-AGG | NHEJ |
| 2-cell | donor A | embryo 7 | nd | nd | T | G | T | A | nd | wt | nd | nd | G | G | T | G | na | loss of paternal allele |
| | | embryo 8 (8 cells) | | | | | | | | | | | | | | | | |
| 2-cell | donor A | embryo 8 | normal | f | T | G | T | C | nd | wt | nd | nd | A | A | nd | nd | na | IH-HR |
| 2-cell | donor A | embryo 8 | f | nd | T | G | T | A | A (100%) | wt | wt (100%) | nd | G | G | T | G | na | loss of paternal allele |
| 2-cell | donor A | embryo 8 | nd | nd | T | G | T | A | A (100%) | wt | wt (100%) | nd | nd | nd | T | G | na | loss of paternal allele |
| 2-cell | donor A | embryo 8 | nd | nd | T | G | T | A | nd | wt | nd | nd | nd | nd | nd | nd | na | loss of paternal allele |
| 2-cell | donor A | embryo 8 | nd | nd | T | G | T | A | nd | wt | nd | nd | G | G | nd | nd | na | loss of paternal allele |
| 2-cell | donor A | embryo 8 | nd | nd | T | G | T | A/C | nd | wt/indel | nd | nd | G/A | G/A | T/- | G/A | TCTGT(-1bp)CTG-AGG | NHEJ |
| 2-cell | donor A | embryo 8 | nd | nd | T | G | T | A | nd | wt | nd | nd | G | G | T | G | na | loss of paternal allele |
| | | embryo (3 of 4 cells/fragments) | | | | | | | | | | | | | | | | |
| 2-cell | donor A | embryo 1 | paternal 6p fragment only | A | T | G | T | f | na | f | na | no detection | f | f | f | f | no allele detected | cleaved chromosome fragment |
| 2-cell | donor A | embryo 1 | del 6q (pat) | G/A | T | G | T | f | na | f | na | no detection | f | f | T | G | no allele detected | no allele detected |
| 2-cell | donor A | embryo 1 | del 6p (pat) paternal, dup 6q (pat) | G | C | A | C | A | A (100%) | wt | wt (100%) | wt/not detected | G | G | T | G | na | loss of paternal allele |
| | | embryo (3 cells) | | | | | | | | | | | | | | | | |
| 2-cell | donor A | embryo 2 | f | f | C | A | C | A | A (100%) | wt | wt (100%) | wt/not detected | nd | nd | T | G | na | loss of paternal allele |
| 2-cell | donor A | embryo 2 | monosomy 6 (pat) | G | C | A | C | A | A (100%) | wt | wt (100%) | wt/not detected | G | A | T | G | na | loss of paternal allele |
| 2-cell | donor A | embryo 2 | del 6q (pat) | G/A | C/T | A/G | C/T | A | A (100%) | wt | wt (100%) | wt/not detected | G | G | T | G | na | telomeric loss of paternal allele |
| 2-cell | donor A | morula multiple cells embryo 10 biopsy 1 | nd | nd | T | G | T | A | A (99%) / C (1%) | wt (mosaic) | wt (99%) / edited (1%) | na | nd | nd | T | G | AGAAAGATCTTTCT(-8bp)G-AGG | MMEJ -8bp |
| 2-cell | donor A | morula multiple cells embryo 10 biopsy 2 | f | nd | T | G | T | A | A (100%) | wt | wt (100%) | na | nd | nd | nd | nd | na | loss of paternal allele |
| 2-cell | donor A | morula multiple cells duplicate embryo 11 | nd | nd | C/T | A/G | C/T | A | A (100%) | wt | wt (100%) | na | nd | nd | T/- | G/A | na | loss of paternal allele |
| na | donor E | gDNA egg donor E | nd | nd | T/T | G/G | T/T | A/C | nd | wt/wt | nd | nd | G/A | G/A | - | A/A | GTCTTTCTTCTGTACTGGAGG | original maternal allele |
| 2-cell | donor E | morula multiple cells embryo 12 | nd | nd | T | G | T | C/C | C (100%) | wt/indel | wt (33%) / edited (67%) | nd | nd | nd | - | A | CTT(-5bp)CTG-AGG | MMEJ -5bp |
| 2-cell | donor E | blastocyst multiple cells embryo 13 | nd | nd | T | G | T | C/C | C (100%) | wt/indel | wt (37%) / edited (63%) | nd | nd | nd | - | A | CTT(-5bp)CTG-AGG | MMEJ -5bp |
| na | donor H | gDNA egg donor H | nd | nd | C/C | A/A | C/C | A/A | nd | wt/wt | nd | nd | G/G | T/T | T/T | G/G | GTCTTTCTTCTGTACTGGAGG | original maternal allele |
| | | embryo 5 (8 cells) | | | | | | | | | | | | | | | | |
| 2-cell | donor H | embryo 5 | nd | nd | C | A | C | A | nd | wt | nd | nd | G | G | T | G | na | loss of paternal allele |
| 2-cell | donor H | embryo 5 | nd | nd | C/T | A/G | C/T | A | A (100%) | wt | wt (100%) | nd | G | G | T/- | G/A | na | loss of paternal allele |
| 2-cell | donor H | embryo 5 | nd | nd | C | A | C | A | nd | wt | nd | nd | G | G | T | G | na | loss of paternal allele |
| 2-cell | donor H | embryo 5 | nd | nd | C/T | A/G | C/T | A/C | nd | wt/fs | nd | nd | f | f | T/- | G/A | na | no change |
| 2-cell | donor H | embryo 5 | nd | nd | T | G | T | A/C | nd | wt/fs | nd | nd | G/A | G/A | T/- | G/A | na | no change |
| 2-cell | donor H | embryo 5 | nd | nd | nd | nd | nd | C | nd | wt | nd | nd | A | A | T/- | G/A | na | IH-HR |
| | | embryo 6 (6 cells) | | | | | | | | | | | | | | | | |
| 2-cell | donor H | embryo 6 | nd | nd | T | G | T | A/C | nd | wt/indel | nd | nd | A | A | T/- | G/A | CTTC(-4bp)CTG-AGG | NHEJ |
| 2-cell | donor H | embryo 6 | nd | nd | T | G | T | A/C | nd | wt/indel | nd | nd | G/A | G/A | T/- | G/A | CTTC(-4bp)CTG-AGG | NHEJ |
| 2-cell | donor H | embryo 6 | nd | nd | C/T | A/G | C/T | A | nd | wt | nd | nd | G | G | T/- | G/A | na | loss of paternal allele |
| 2-cell | donor H | embryo 6 | nd | nd | C | A | C | A | nd | wt | nd | nd | G | G | T | G | na | loss of paternal allele |
| 2-cell | donor H | embryo 6 | nd | nd | C | A | C | A/A | nd | wt/indel | nd | nd | G/G | G/G | T | G | CTGTA ACTGGAGG | loss of paternal allele, wt maternal/altered maternal allele NHEJ |
| | | embryo 9 (10 cells) | | | | | | | | | | | | | | | | |
| 2-cell | donor H | embryo 9 | nd | nd | C | A | C | A | nd | wt | nd | nd | G | G | T | G | na | loss of paternal allele |
| 2-cell | donor H | embryo 9 | nd | nd | C | A | C | A | nd | wt | nd | nd | G | G | T/- | G/A | na | loss of paternal allele |
| 2-cell | donor H | embryo 9 | nd | nd | C/T | A/G | C/T | A/C | A (53%) / C (47%) | wt/indel | wt (53%) / edited (47%) | nd | G/A | G/A | T/- | G/A | AGAAAGATCTTTCT(-8bp)G-AGG | MMEJ -8bp |
| 2-cell | donor H | embryo 9 | nd | nd | C/T | A/G | C/T | A | nd | wt/indel | nd | nd | G/A | A | T/- | G/A | CTT(-5bp)CTG-AGG | MMEJ -5bp |
| 2-cell | donor H | embryo 9 | nd | nd | C/T | A/G | C/T | A | A (100%) | wt | wt (100%) | nd | G | G | T | G | na | telomeric loss of paternal allele |
| 2-cell | donor H | embryo 9 | nd | nd | C/T | A/G | C/T | A/C | nd | wt/indel | nd | nd | G/A | G/A | nd | G/A | CTT(-5bp)CTG-AGG | MMEJ -5bp |
| 2-cell | donor H | embryo 9 | nd | nd | T | G | T | C | C (100%) | wt | wt (100%) | nd | A | A | nd | nd | na | IH-HR |
| 2-cell | donor H | embryo 9 | nd | nd | C | A | C | A | nd | wt | nd | nd | G | G | T | G | na | loss of paternal allele |
| **other events** | | | | | | | | | | | | | | | | | | |
| na | na | gDNA sperm donor | normal | A/A | T/T | G/G | T/T | C/C | C/C | fs/fs | fs/fs | fs/fs | A/A | A/A | -/- | A/A | GTCTTTCTTCTGTACTG-AGG | original paternal allele |
| na | donor A | gDNA egg donor A | normal | G/G | C/T | A/G | C/T | A/A | nd | wt/wt | nd | wt/wt | G/G | G/G | T/T | G/G | GTCTTTCTTCTGTACTGGAGG | original maternal allele |
| 2PN | donor A | blastomere cell (2 cells) | normal | G/A | C/T | A/G | C/T | A/C | A(32%) / C (68%) | wt/indel | wt (33%) / edited (67%) | wt/not detected | G/A | G/A | T/- | G/A | TCTTCTGT(-1bp)CTG-AGG | NHEJ |
| 2PN | donor A | sister blastomere cell (2 cells) | normal | f | nd | nd | nd | A | nd | wt | nd | wt/not detected | nd | nd | nd | nd | na | loss of paternal allele |
| na | donor G | gDNA egg donor G | normal | nd | C/C | A/A | C/C | A/A | A/A | wt/wt | nd | nd | G/G | G/G | T/T | G/G | GTCTTTCTTCTGTACTGGAGG | original maternal allele |
| 2-cell androgenesis | donor G | blastomere androgenesis | nd | A | T | G | T | C | nd | indel | nd | nd | A | A | T/- | G/A | CTT(-5bp)CTG-AGG | MMEJ -5bp androgenesis |

### Table S2

| | donor egg genome origin | Sanger genotype | Target Reads | Read Errors | Significant Reads | WT Reads | Mutant ($EYS^{2265fs}$) Reads | paternal edited Reads | % wt | % $EYS^{2265fs}$ | % edited | Bases (reverse orientation from Figures) | RS66502009 maternal | RS66502009 paternal | % paternal SNP |
|---|---|---|---|---|---|---|---|---|---|---|---|---|---|---|---|
| **ES cell line experiments, gRNA specificity (relates to Fig. 1D & 4D)** | | | | | | | | | | | | | | | |
| eysES6 & Cas9 gRNA | sperm/donor F | mosaic edited | 38241 | 0 | 38241 | 18949 | 6508 | 12784 | 49.55% | 17.02% | 33.43% | mosaic | 19493 | 18642 | 48.9 |
| eysES6 & Cas9 gRNA | sperm/donor F | mosaic edited | 39154 | 0 | 39154 | 18697 | 17934 | 2523 | 47.75% | 45.80% | 6.44% | mosaic | 19457 | 19697 | 50.3 |
| Control eysES6, no Cas9 | sperm/donor F | wt/ *EYS* $^{2265fs}$ | 41128 | 0 | 41128 | 19608 | 21520 | - | 47.68% | 52.32% | 0.00% | CAGATGGAAAACCT-CAGTACAGAAGA | 20022 | 21106 | 51.3 |
| **MII injection (relates to Fig. 2C)** | | | | | | | | | | | | | | | |
| single oocyte control | donor G | haploid WT | 77525 | 211 | 77314 | 77314 | - | - | 100.00% | 0.00% | 0.00% | CAGATGGAAAACCTCCAGTACAGAAGA | 76939 | - | 0.0 |
| sperm control | EYS | haploid mutant | 48245 | 135 | 48110 | - | 48110 | - | 0.00% | 100.00% | 0.00% | CAGATGGAAAACCT-CAGTACAGAAGA | - | 48245 | 100.0 |
| zygote 1 @ 20h | donor A | WT | 83382 | 275 | 83107 | 83107 | - | - | 100.00% | 0.00% | 0.00% | CAGATGGAAAACCTCCAGTACAGAAGA | 82778 | - | 0.0 |
| zygote 2 @ 20h | donor A | WT | 51826 | 311 | 51515 | 51515 | - | - | 100.00% | 0.00% | 0.00% | CAGATGGAAAACCTCCAGTACAGAAGA | 51608 | - | 0.0 |
| blastomere embryo 1 | donor A | WT | 92864 | 231 | 92633 | 92633 | - | - | 100.00% | 0.00% | 0.00% | CAGATGGAAAACCTCCAGTACAGAAGA | 92147 | - | 0.0 |
| blastomere embryo 1 | donor A | WT | 56495 | 110 | 56385 | 56385 | - | - | 100.00% | 0.00% | 0.00% | CAGATGGAAAACCTCCAGTACAGAAGA | 51133 | 5051 | 9.0 |
| blastomere embryo 1 | donor A | WT | 52752 | 136 | 52616 | 52616 | - | - | 100.00% | 0.00% | 0.00% | CAGATGGAAAACCTCCAGTACAGAAGA | 52399 | - | 0.0 |
| blastomere embryo 1 | donor A | WT | 53876 | 160 | 53716 | 53716 | - | - | 100.00% | 0.00% | 0.00% | CAGATGGAAAACCTCCAGTACAGAAGA | 53504 | - | 0.0 |
| blastomere embryo 1 | donor A | WT | 54989 | 149 | 54840 | 54840 | - | - | 100.00% | 0.00% | 0.00% | CAGATGGAAAACCTCCAGTACAGAAGA | 54634 | - | 0.0 |
| blastomere embryo A | donor D | WT/indel | 34498 | 0 | 34498 | 23224 | 0 | 11274 | 67.32% | 0.00% | 32.68% | CAGATGGAAAACCT-CAG--AAGAAGAA | na | na | na |
| blastomere embryo A | donor D | WT/indel | 28946 | 109 | 28837 | 16095 | 0 | 12742 | 55.81% | 0.00% | 44.19% | CAGATGGAAAACCT-CAG--AAGAAGAA | na | na | na |
| blastomere embryo A | donor D | WT/indel | 50609 | 0 | 50609 | 22627 | - | 27982 | 44.71% | 0.00% | 55.29% | CAGATGGAAAACCT-CAG--AAGAAGAAAGA | na | na | na |
| blastomere embryo A | donor D | WT/indel | 31471 | 0 | 31471 | 25233 | 0 | 6238 | 80.18% | 0.00% | 19.82% | CAGATGGAAAACCT-CAG--AAGAAGAA | na | na | na |
| blastomere embryo A | donor D | WT | 59985 | 168 | 59817 | 59817 | - | - | 100.00% | 0.00% | 0.00% | CAGATGGAAAACCTCCAGTACAGAAGA | na | na | na |
| blastomere embryo B | donor D | WT/indel | 35352 | 0 | 35352 | 32543 | 0 | 2809 | 92.05% | 0.00% | 7.95% | CAGATGGAAAACCT------CAGAAGAAA | na | na | na |
| blastomere embryo B | donor D | WT/indel | 31096 | 0 | 31096 | 7130 | 0 | 23966 | 22.93% | 0.00% | 77.07% | CAGATGGAAAACCT------CAGAAGAAA | na | na | na |
| blastomere embryo B | donor D | WT/indel | 36255 | 0 | 36255 | 11682 | 0 | 24573 | 32.22% | 0.00% | 67.78% | CAGATGGAAAACCT------CAGAAGAAA | na | na | na |
| blastomere embryo B | donor D | WT/indel | 31046 | 0 | 31046 | 24516 | 0 | 6530 | 78.97% | 0.00% | 21.03% | CAGATGGAAAACCT------CAGAAGAAA | na | na | na |
| blastomere embryo C | donor D | WT | 35278 | 0 | 35278 | 35278 | 0 | 0 | 100.00% | 0.00% | 0.00% | CAGATGGAAAACCTCCAGTACAGAAGA | na | na | na |
| blastomere embryo C | donor D | WT | 31679 | 0 | 31679 | 31679 | 0 | 0 | 100.00% | 0.00% | 0.00% | CAGATGGAAAACCTCCAGTACAGAAGA | na | na | na |
| blastomere embryo C | donor D | WT | 29768 | 0 | 29768 | 29768 | 0 | 0 | 100.00% | 0.00% | 0.00% | CAGATGGAAAACCTCCAGTACAGAAGA | na | na | na |
| blastomere embryo C | donor D | WT | 37798 | 121 | 37677 | 37677 | 0 | 0 | 100.00% | 0.00% | 0.00% | CAGATGGAAAACCTCCAGTACAGAAGA | na | na | na |
| TE biopsy d6 (embryo E) | donor D | WT | 46779 | 133 | 46646 | 46646 | - | - | 100.00% | 0.00% | 0.00% | CAGATGGAAAACCTCCAGTACAGAAGA | na | na | na |
| TE biopsy d6 (embryo F) | donor D | WT | 52020 | 129 | 51891 | 51891 | - | - | 100.00% | 0.00% | 0.00% | CAGATGGAAAACCTCCAGTACAGAAGA | na | na | na |
| **2-cell injections (relates to Fig. 3D)** | | | | | | | | | | | | | | | |
| 2-cell inj. embryo 8 | donor A | WT | 41454 | 0 | 41454 | 41454 | - | - | 100.00% | 0.00% | 0.00% | CAGATGGAAAACCTCCAGTACAGAAGA | 41246 | - | 0.0 |
| 2-cell inj. embryo 8 | donor A | WT | 61682 | 168 | 61514 | 61514 | - | - | 100.00% | 0.00% | 0.00% | CAGATGGAAAACCTCCAGTACAGAAGA | 61380 | - | 0.0 |
| 2-cell inj blastomere embryo 1 | donor A | WT | 47822 | 100 | 47722 | 47722 | - | - | 100.00% | 0.00% | 0.00% | CAGATGGAAAACCTCCAGTACAGAAGA | 47695 | - | 0.0 |
| 2-cell inj blastomere embryo 2 | donor A | WT | 44403 | 0 | 44403 | 44403 | - | - | 100.00% | 0.00% | 0.00% | CAGATGGAAAACCTCCAGTACAGAAGA | 44287 | - | 0.0 |
| 2-cell inj blastomere embryo 2 | donor A | WT | 47985 | 0 | 47985 | 47985 | - | - | 100.00% | 0.00% | 0.00% | CAGATGGAAAACCTCCAGTACAGAAGA | 47985 | - | 0.0 |
| 2-cell inj blastomere embryo 2 | donor A | WT | 55562 | 118 | 55444 | 55444 | - | - | 100.00% | 0.00% | 0.00% | CAGATGGAAAACCTCCAGTACAGAAGA | 55431 | - | 0.0 |
| 2-cell inj. Embryo 5 | donor H | WT | 56398 | 165 | 56233 | 56233 | - | - | 100.00% | 0.00% | 0.00% | CAGATGGAAAACCTCCAGTACAGAAGA | 56012 | - | 0.0 |
| 2-cell in. embryo 9 | donor H | Homozygous WT, IH-HR | 49564 | 147 | 49417 | 49417 | - | - | 100.00% | 0.00% | 0.00% | CAGATGGAAAACCTCCAGTACAGAAGA | - | 49564 | 100.0 |
| 2-cell in. embryo 9 | donor H | Heterozygous WT, MMEJ | 46458 | 0 | 46458 | 24518 | - | 21940 | 52.77% | 0.00% | 47.23% | CAGATGGAAAACCTC---------AGAAAGA | 23685 | 20987 | 47.0 |
| 2-cell in. embryo 9 | donor H | WT | 54588 | 191 | 54397 | 54397 | - | - | 100.00% | 0.00% | 0.00% | CAGATGGAAAACCTCCAGTACAGAAGA | 54303 | - | 0.0 |
| morula multiple cells embryo 10 biopsy 1 | donor A | mosaic WT and MMEJ | 48819 | 101 | 48718 | 48335 | - | 383 | 99.21% | 0.00% | 0.79% | CAGATGGAAAACCTC---------AGAAAGAC | 48021 | 638 | 1.3 |
| morula multiple cells embryo 10 biopsy 2 | donor A | Homozygous WT | 44947 | 119 | 44828 | 44828 | - | - | 100.00% | 0.00% | 0.00% | CAGATGGAAAACCTCCAGTACAGAAGA | 44634 | - | 0.0 |
| blastocyst multiple cells embryo 13 | donor E | MMEJ/WT | 55221 | 100 | 55121 | 20487 | - | 34634 | 37.17% | 0.00% | 62.83% | CAGATGGAAAACCT------CAGAAGAAAGA | na | na | na |
| morula multiple cells duplicate embryo 11 | donor A | Homozygous WT | 54758 | 139 | 54619 | 54619 | - | - | 100.00% | 0.00% | 0.00% | CAGATGGAAAACCTCCAGTACAGAAGA | 54624 | - | 0.0 |
| morula multiple cells embryo 12 | donor E | MMEJ/WT | 43145 | 0 | 43145 | 13991 | - | 29154 | 32.43% | 0.00% | 67.57% | CAGATGGAAAACCT------CAGAAGAAA | na | na | na |
| **other events** | | | | | | | | | | | | | | | |
| blastomere from 2PN injection NHEJ | donor A | NHEJ/WT | 53681 | 0 | 53681 | 17513 | - | 36168 | 32.62% | 0.00% | 67.38% | CAGATGGAAAACCT-CAG-ACAGAAGA | 16778 | 36903 | 68.7 |

### Table S3

| sample name | egg donor | rs758109813 genotype call (Sanger sequencing) | chromosome 6 | karyotype | display item | breakpoint calling | chromosome 1 | chromosome 2 | chromosome 3 | chromosome 4 | chromosome 5 | chromosome 6 | chromosome 7 | chromosome 8 | chromosome 9 | chromosome 10 | chromosome 11 | chromosome 12 | chromosome 13 | chromosome 14 | chromosome 15 | chromosome 16 | chromosome 17 | chromosome 18 | chromosome 19 | chromosome 20 | chromosome 21 | chromosome 22 |
|---|---|---|---|---|---|---|---|---|---|---|---|---|---|---|---|---|---|---|---|---|---|---|---|---|---|---|---|---|
| **Karyotypes of pluripotent stem cells** | | | | | | | | | | | | | | | | | | | | | | | | | | | | |
| eysES4 | donor C | wt/indel | normal | 46, XX | Fig. S1B | na | normal | normal | normal | normal | normal | **normal** | normal | normal | normal | normal | normal | normal | normal | normal | normal | normal | normal | normal | normal | normal | normal | normal |
| eysES5 | donor C | wt/indel | normal | 46, XY | Fig. S1B | na | normal | normal | normal | normal | normal | **normal** | normal | normal | normal | normal | normal | normal | normal | normal | normal | normal | normal | normal | normal | normal | normal | normal |
| eysES6 | donor C | wt/fs (no change) | normal | 46, XY | Fig. S1B | na | normal | normal | normal | normal | normal | **normal** | normal | normal | normal | normal | normal | normal | normal | normal | normal | normal | normal | normal | normal | normal | normal | normal |
| eysES8 | donor C | wt/indel | normal | 46, XY | Fig. S1B | na | normal | normal | normal | normal | normal | **normal** | normal | normal | normal | normal | normal | normal | normal | normal | normal | normal | normal | normal | normal | normal | normal | normal |
| single eysES4 cell | donor C | wt/indel | normal | 46, XX | Fig. S1B | na | normal | normal | normal | normal | normal | **normal** | normal | normal | normal | normal | normal | normal | normal | normal | normal | normal | normal | normal | normal | normal | normal | normal |
| single eysES4 cell | donor C | wt/indel | normal | 46, XX | Fig. S1B | na | normal | normal | normal | normal | normal | **normal** | normal | normal | normal | normal | normal | normal | normal | normal | normal | normal | normal | normal | normal | normal | normal | normal |
| single control eysES6 cell | donor C | wt/fs (no change) | normal | 46, XY | Fig. S1B | na | normal | normal | normal | normal | normal | **normal** | normal | normal | normal | normal | normal | normal | normal | normal | normal | normal | normal | normal | normal | normal | normal | normal |
| single control eysES6 cell | donor C | wt/fs (no change) | normal | 46, XY | na | na | normal | normal | normal | normal | normal | **normal** | normal | normal | normal | normal | normal | normal | normal | normal | normal | normal | normal | normal | normal | normal | normal | normal |
| **Karyotypes after MII injection** | | | | | | | | | | | | | | | | | | | | | | | | | | | | |
| TE biopsy embryo 4 (corresponds to eysES4) | donor C | wt/indel | normal | 46, XX | Fig. S1A | na | normal | normal | normal | normal | normal | **normal** | normal | normal | normal | normal | normal | normal | normal | normal | normal | normal | normal | normal | normal | normal | normal | normal |
| TE biopsy embryo 5 (corresponds to eysES5) | donor C | wt/indel | normal | 46, XY | Fig. S1A | na | normal | normal | normal | normal | normal | **normal** | normal | normal | normal | normal | normal | normal | normal | normal | normal | normal | normal | normal | normal | normal | normal | normal |
| TE biopsy embryo 6 (corresponds to eysES6) | donor C | wt/fs (no change) | normal | 46, XY | Fig. S1A | na | normal | normal | normal | normal | normal | **normal** | normal | normal | normal | normal | normal | normal | normal | normal | normal | normal | normal | normal | normal | normal | normal | normal |
| TE biopsy embryo 8 (corresponds to eysES8) | donor C | wt/indel | normal | 46, XY | Fig. S1A | na | normal | normal | normal | normal | normal | **normal** | normal | normal | normal | normal | normal | normal | normal | normal | normal | normal | normal | normal | normal | normal | normal | normal |
| MII injection TE BIOPSY embryo F | donor D | wild type | monosomy 6 | 45,XY,-6 | Fig. 2 | nd | normal | normal | normal | normal | normal | **monosomy 6 (pat)** | normal | normal | normal | normal | normal | normal | normal | normal | normal | normal | normal | normal | normal | normal | normal | normal |
| MII injection TE BIOPSY embryo E | donor D | wild type | del(6q) (pat) | 46,XY,del(6q) breakpoint near EYS | Fig. 2 | <101kb from mutation site | normal | normal | normal | normal | normal | **del 6q (pat)** | normal | normal | normal | normal | normal | normal | normal | normal | normal | normal | normal | normal | normal | normal | normal | normal |
| MII injection TE BIOPSY embryo D | donor D | wt/indel | normal | 46,XX | Fig. 2 | na | normal | normal | normal | normal | normal | **normal** | normal | normal | normal | normal | normal | normal | normal | normal | normal | normal | normal | normal | normal | normal | normal | normal |
| MII injection TE biopsy embryo 10 | donor C | nd | normal | 46, XX | Fig. S1A | na | normal | normal | normal | normal | normal | **normal** | normal | normal | normal | normal | normal | normal | normal | normal | normal | normal | normal | normal | normal | normal | normal | normal |
| MII injection blastomere embryo B1 | donor D | wt/indel | normal | del 12p (pat) | Fig. S2D | na | normal | normal | normal | normal | normal | **normal** | normal | normal | normal | normal | normal | del12p (pat)* | normal | normal | normal | normal | normal | normal | normal | normal | normal | normal |
| MII injection blastomere embryo B1 | donor D | wt/indel | normal | del 12p (pat) | Fig. S2D | na | normal | normal | normal | normal | normal | **normal** | normal | normal | normal | normal | normal | del12p (pat) * | normal | normal | normal | normal | normal | normal | normal | normal | normal | normal |
| MII injection blastomere embryo B1 | donor D | wt/indel | normal | del 12p (pat) | Fig. S2D | na | normal | normal | normal | normal | normal | **normal** | del7q (pat) | normal | normal | normal | normal | del12p (pat) * | normal | normal | normal | normal | normal | normal | normal | normal | normal | normal |
| MII injection blastomere embryo B1 | donor D | wt/indel | normal | del 12p (pat) | Fig. S2D | na | normal | normal | normal | normal | normal | **normal** | normal | normal | normal | normal | normal | del12p (pat) * | normal | normal | normal | normal | normal | normal | normal | normal | normal | normal |
| MII injection single 'wt' blastomere embryo C | donor D | wild type | del(6) (pat) | chaotic aneuploid | Fig. S2A | na | normal | normal | normal | normal | monosomy 5 (pat) | **monosomy 6 (pat)** | normal | normal | normal | monosomy 10 (pat) | normal | monosomy 12 (pat) | monosomy 13 (pat) | normal | normal | normal | normal | normal | monosomy 19 (pat) | normal | normal | normal |
| MII injection single 'wt' blastomere embryo C | donor D | wild type | del(6) (pat) | chaotic aneuploid | Fig. S2A | na | monosomy 1 (mat) | monosomy 2 (mat) | normal | monosomy 4 (mat) | monosomy 5 (pat) | **monosomy 6 (pat)** | monosomy 7 (mat) | monosomy 8 (mat) | normal | monosomy 10 (pat) | normal | normal | normal | normal | normal | normal | monosomy 17 (mat) | normal | monosomy 19 (pat) | monosomy 20 (mat) | normal | monosomy 22 (mat) |
| MII injection single 'wt' blastomere embryo C | donor D | wild type | del(6) (pat) | chaotic aneuploid | Fig. S2A | na | normal | normal | normal | normal | monosomy 5 (pat) | **monosomy 6 (pat)** | normal | del8p23.1 (mat) | normal | monosomy 10 (pat) | normal | normal | normal | normal | normal | normal | normal | normal | monosomy 19 (pat) | normal | normal | normal |
| MII injection single 'wt' blastomere embryo C | donor D | wild type | del(6) (pat) | chaotic aneuploid | Fig. S2A | na | monosomy 1 (mat) | monosomy 2 (mat) | normal | monosomy 4 (mat) | monosomy 5 (pat) | **monosomy 6 (pat)** | monosomy 7 (mat) | monosomy 8 (mat) | normal | monosomy 10 (pat) | normal | normal | normal | normal | normal | normal | monosomy 17 (mat) | normal | monosomy 19 (pat) | monosomy 20 (mat) | normal | monosomy 22 (mat) |
| MII injection single blastomere embryo C | donor D | no allele | paternal 6p only | chaotic aneuploid | Fig. S2A | <573Kb | monosomy 1 (pat) | normal | nullisomy 3 | monosomy 4 (pat) | normal | **paternal 6p only** | normal | nullisomic | monosomy 9 (mat) | normal | nullisomic | normal | normal | normal | nullisomic | normal | monosomy 17 (pat) | monosomy 18 (mat) | nullisomy 19 | monosomy 20 (pat) | monosomy 21 (pat) | normal |
| MII injection single blastomere embryo C | donor D | no allele | paternal 6p only | chaotic aneuploid | Fig. S2A | <573Kb | monosomy 1 (pat) | normal | nullisomy 3 | monosomy 4 (pat) | normal | **paternal 6p only** | normal | nullisomic | monosomy 9 (mat) | normal | nullisomic | normal | normal | normal | nullisomic | normal | monosomy 17 (pat) | monosomy 18 (mat) | normal | monosomy 20 (pat) | monosomy 21 (pat) | normal |
| MII injection single 'wt' blastomere embryo C | donor D | indel (mat) | dup(6q)(pat) | chaotic aneuploid | Fig. S2A | <573Kb | normal | monosomy 2 (pat) | normal | normal | monosomy 5 (mat) | **dup(6q)(pat)** | monosomy 7 (pat) | complex rearrangements | monosomy 9 (pat) | monosomy 10 (mat) | normal | normal | normal | nullisomic | normal | nullisomic | normal | monosomy 18 (pat) | monosomy 19 (mat) | normal | monosomy 21 (mat) | monosomy 22 (pat) |
| MII injection single 'wt' blastomere embryo 1 | donor A | wild type | failed | failed | na | na | no call | no call | no call | no call | no call | no call | no call | no call | no call | no call | no call | no call | no call | no call | no call | no call | no call | no call | no call | no call | no call | no call |
| MII injection single 'wt' blastomere embryo 1 | donor A | wild type | del 6q (pat) | normal for other chromosomes | Fig. S2B | <573Kb | normal | normal | normal | normal | normal | **del(6q) (pat)** | normal | normal | normal | normal | normal | normal | normal | normal | normal | normal | normal | normal | normal | normal | normal | normal |
| MII injection single 'wt' blastomere embryo 1 | donor A | wild type | del 6q (pat) | normal for other chromosomes | Fig. S2B | <573Kb | normal | normal | normal | normal | normal | **del(6q)(pat)** | normal | normal | normal | normal | normal | normal | normal | normal | normal | normal | normal | normal | normal | normal | normal | normal |
| MII injection single 'wt' blastomere embryo 1 | donor A | wild type | dup 6q (pat) | 45,XY,-5,-13,+15 | Fig. S2B | <573Kb | normal | normal | normal | normal | monosomy 5 (pat) | **dup (6q)(pat)** | normal | normal | normal | normal | normal | normal | monosomy 13 (pat) | normal | trisomy 15 (mat) | normal | normal | normal | normal | normal | normal | normal |
| MII injection single 'wt' blastomere embryo 1 | donor A | wild type | del 6q (pat) | monosomy 5 | Fig. S2B | <573Kb | normal | normal | normal | normal | monosomy 5 (pat) | **del(6q) (pat)** | normal | normal | normal | normal | normal | normal | normal | normal | normal | normal | normal | normal | normal | normal | normal | normal |
| MII injection blastomere embryo A | donor D | wt/indel | maternal del del6q21-27 (mat) | chaotic aneuploid | Fig. S2C | na | normal | monosomy 2 (pat) | normal | normal | del 5q32-tel (mat) | **del (6q21-tel) (mat) (Fra6F)** | monosomy 7 (mat) | normal | monosomy 9 (mat) | normal | monosomy 11 (pat) | normal | normal | normal | monosomy 15 (pat) | monosomy 16q (pat) | monosomy 17 (mat) | normal | monosomy 19 (pat) | normal | normal | normal |
| MII injection blastomere embryo A | donor D | wt/indel | normal | normal | Fig. S2C | na | normal | normal | normal | normal | normal | **normal** | normal | normal | normal | normal | normal | normal | normal | normal | normal | normal | normal | normal | normal | normal | normal | normal |
| MII injection blastomere embryo A | donor D | wt/indel | normal | chaotic aneuploid | Fig. S2C | na | del 1q25.2-tel (pat) | monosomy 2 (pat) | normal | normal | normal | **normal** | monosomy 7 (mat) | normal | monosomy 9 (mat) | normal | monosomy 11 (pat) | normal | normal | normal | monosomy 15 (pat) | normal | monosomy 17 (mat) | normal | monosomy 19 (pat) | normal | normal | normal |
| MII injection blastomere embryo A | donor D | wt/indel | normal | normal | Fig. S2C | na | normal | normal | normal | normal | normal | **normal** | normal | normal | normal | normal | normal | normal | normal | normal | normal | normal | normal | normal | normal | normal | normal | normal |
| MII injection blastomere embryo A8 | donor D | wild type | monosomy 6 (pat) | chaotic aneuploid | Fig. S2C | na | nullisomy 1 | monosomy 2 (pat) | monosomy 3 (mat) | nullisomic | nullisomic | **monosomy 6 (pat)** | nullisomic | monosomy 8 (mat) | monosomy 9 (mat) | monosomy 10 (pat) | nullisomic | monosomy 12 (mat) | nullisomic | monosomy 14 (mat) | nullisomic | monosomy 16 (pat) | monosomy 17 (mat) | monosomy 18 (mat) | nullisomic | monosomy 20 (pat) | nullisomic | nullisomic |
| MII injection 'wt' zygote 1 | donor A | wild type | normal | 46,XX | Fig. 2F | none | normal | normal | normal | normal | normal | **normal** | normal | normal | normal | normal | normal | normal | normal | normal | normal | normal | normal | normal | normal | normal | normal | normal |
| MII injection 'wt' zygote 2 | donor A | wild type | normal | normal | na | na | normal | normal | normal | normal | normal | **normal** | normal | normal | normal | normal | normal | normal | normal | normal | normal | normal | normal | normal | normal | normal | normal | normal |
| **Karyotypes after 2-cell injections** | | | | | | | | | | | | | | | | | | | | | | | | | | | | |
| 2-cell injection, blastomere embryo 1 | donor A | no allele detected | paternal 6p only | chr6p fragment only | Fig. 3 | nd | nullisomic | nullisomic | nullisomic | nullisomic | nullisomic | **paternal 6p only** | nullisomic | nullisomic | nullisomic | nullisomic | nullisomic | nullisomic | nullisomic | nullisomic | nullisomic | nullisomic | nullisomic | nullisomic | nullisomic | nullisomic | nullisomic | nullisomic |
| 2-cell injection, blastomere embryo 1 | donor A | no allele detected | del(6q)(pat) | del 16q22-tel (mat), del18p (mat) | Fig. 3 | <101kb from mutation site | normal | normal | normal | normal | normal | **del(6q)(pat)** | normal | normal | normal | normal | normal | normal | normal | normal | normal | del 16q22-tel (mat) | normal | del18p (mat) FRA18E | normal | normal | normal | normal |
| 2-cell injection, blastomere embryo 1 | donor A | wild type | del(6p)(pat),dup(6q)(pat) | del 16q22-tel (pat), del18p (pat) | Fig. 3 | nd | normal | normal | normal | normal | normal | **del(6p)(pat),dup(6q)(pat)** | normal | normal | normal | normal | normal | normal | normal | normal | normal | del 16q22-tel (mat) | normal | dup18p (mat) FRA18E | normal | normal | normal | normal |
| 2-cell injection, blastomere embryo 2 | donor A | wild type | failed | chaotic aneuploidy | Fig. S3 | na | inconclusive/low signal | monosomy 2 (pat) | normal | normal | normal | inconclusive/low signal | normal | del 8q23.2-23.3 (pat) | monosomy 9 (pat) | normal | monosomy 11 (mat) | normal | monosomy 13 (pat) | normal | normal | monosomy 16 (mat) | inconclusive/low signal | normal | normal | inconclusive/low signal | monosomy 21 (pat) | normal |
| 2-cell injection, blastomere embryo 2 | donor A | wild type | monosomy 6 (pat) | chaotic aneuploidy | Fig. S3 | na | monosomy 1 (mat) | monosomy 2 (pat) | monosomy 3 (mat) | normal | monosomy 5 (pat) | **monosomy 6 (pat)** | normal | normal | monosomy 9 (mat) | normal | monosomy 11 (pat) | monosomy 12 (mat) | monosomy 13 (mat) | normal | monosomy 15 (mat) | normal | normal | monosomy 18 (pat) | normal | normal | monosomy 21 (pat) | normal |
| 2-cell injection, blastomere embryo 2 | donor A | wild type | del(6q)(pat) | chaotic aneuploidy | Fig. S3 | <-10kb of mutation site | normal | monosomy 2 (mat) | normal | monosomy 4 (pat) | monosomy 5 (mat) | **del(6q)(pat)** | normal | normal | normal | monosomy 10 (mat) | monosomy 11 (pat) | normal | monosomy 13 (mat) | normal | normal | monosomy 16 (pat) | normal | normal | normal | normal | monosomy 21 (mat) | normal |
| 2-cell injection, blastomere embryo 9 (possible IH-HR) | donor A | wild type | normal 6 | 46,XXY,-10 | Fig. S3 | na | normal | normal | normal | normal | normal | **normal** | normal | normal | normal | del 10p, (FRA10G) q11.22 (pat), maternal chr10 loss | normal | normal | normal | normal | normal | normal | normal | normal | normal | normal | normal | normal |
| **other events** | | | | | | | | | | | | | | | | | | | | | | | | | | | | |
| 2PN injection | donor A | wild type | normal 6 | normal | na | na | normal | normal | normal | normal | normal | **normal** | normal | normal | normal | normal | normal | normal | normal | normal | normal | normal | normal | normal | normal | normal | normal | normal |
| 2PN injection sibling blastomere | donor A | wt/indel | normal 6 | normal | na | na | normal | normal | normal | normal | normal | **normal** | normal | normal | normal | normal | normal | normal | normal | normal | normal | del 16q22.2 (pat) | normal | normal | normal | normal | normal | normal |

| Summary (excluding nullisomies) | chromosome 1 | chromosome 2 | chromosome 3 | chromosome 4 | chromosome 5 | chromosome 6 | chromosome 7 | chromosome 8 | chromosome 9 | chromosome 10 | chromosome 11 | chromosome 12 | chromosome 13 | chromosome 14 | chromosome 15 | chromosome 16 | chromosome 17 | chromosome 18 | chromosome 19 | chromosome 20 | chromosome 21 | chromosome 22 | sum excluding chr6 | mitotic paternal aneuploidies (excl. chr.6) | mitotic maternal aneuploidies (excl. chr.6) |
|---|---|---|---|---|---|---|---|---|---|---|---|---|---|---|---|---|---|---|---|---|---|---|---|---|---|
| paternal whole chromosome aneuploidies | 2 | 6 | 0 | 3 | 7 | **7** | 1 | 0 | 2 | 5 | 4 | 1 | 3 | 0 | 2 | 2 | 2 | 2 | 6 | 3 | 4 | 1 | 56 (pat) | 62 | 58 |
| maternal whole chromosome aneuploidies | 3 | 3 | 2 | 2 | 2 | **0** | 4 | 3 | 6 | 3 | 1 | 2 | 2 | 1 | 2 | 1 | 5 | 3 | 1 | 2 | 2 | 2 | 52 (mat) | | |
| | | | | | | | | | | | | | | | | | | | | | | | **sum excluding chr.6** | | |
| paternal mitotic segmental | 1 | 0 | 0 | 0 | 0 | **12** | 1 | 1 | 0 | 1 | 0 | 0 | 0 | 0 | 0 | 2 | 0 | 0 | 0 | 0 | 0 | 0 | 6 (pat) | | |
| maternal mitotic segmental | 0 | 0 | 0 | 0 | 1 | **1** | 0 | 1 | 0 | 0 | 0 | 0 | 0 | 0 | 0 | 2 | 0 | 2 | 0 | 0 | 0 | 0 | 6 (mat) | | |
| meiotic loss | 0 | 0 | 0 | 0 | 0 | **0** | 0 | 0 | 0 | 0 | 0 | 4* | 0 | 0 | 0 | 0 | 0 | 0 | 0 | 0 | 0 | 0 | 4 | | |

Table S4

**Color Key**

| |
|---|
| gamete donor |
| 2PN zygote |
| single blastomeres |
| morula (5-15 cells) |
| Trophectoderm (TE) biopsies (5-10 cells) |
| embryonic stem cell |

| time point of Injection of Cas9 RNP | Sample Type | Allele1 Ct | Allele2 Ct | *WT Delta Rn* | *2265fs Delta Rn* | Allele Call |
|---|---|---|---|---|---|---|
| **DONOR GENOMIC DNA** | | | | | | |
| na | genomic DNA sperm donor | 39.523 | 26.812 | 0.257 | 4.103 | homozygous EYS2265fs Allele |
| na | genomic DNA donor B | 26.893 | 30.414 | 2.885 | 0.509 | maternal |
| na | genomic DNA donor C | 26.934 | 30.904 | 2.872 | 0.424 | WT Allele / WT Allele |
| na | genomic DNA donor A | 24.787 | 31.295 | 2.672 | 0.372 | WT Allele / WT Allele |
| na | genomic DNA donor D | 26.429 | 31.351 | 2.672 | 0.334 | WT Allele / WT Allele |
| **EMBRYOS AFTER MII INJECTION of Cas9 RNP** | | | | | | |
| MII | zygote 1 | 32.837 | 37.269 | 1.992 | 0.145 | WT/ altered paternal allele |
| MII | zygote 2 | 32.875 | 34.826 | 1.876 | 0.294 | WT/ altered paternal allele |
| MII | single blastomere embryo 1 | 31.395 | 39.550 | 2.271 | 0.171 | WT/ altered paternal allele |
| MII | single blastomere embryo 1 | 30.794 | 36.367 | 2.139 | 0.201 | WT/ altered paternal allele |
| MII | single blastomere embryo 1 | 32.258 | 35.719 | 1.890 | 0.290 | WT/ altered paternal allele |
| MII | single blastomere embryo A | 31.655 | 38.701 | 2.059 | 0.188 | WT/ altered paternal allele |
| MII | single blastomere embryo A | 31.563 | 37.157 | 2.173 | 0.179 | WT/ altered paternal allele |
| MII | single blastomere embryo A | 27.772 | 33.889 | 2.337 | 0.182 | WT/ altered paternal allele |
| MII | single blastomere embryo A | 28.190 | 31.872 | 2.354 | 0.225 | WT/ altered paternal allele |
| MII | single blastomere embryo A | 28.843 | 33.034 | 2.484 | 0.240 | WT/ altered paternal allele |
| MII | single blastomere embryo B | 24.378 | 30.287 | 1.604 | 0.127 | WT/ altered paternal allele |
| MII | single blastomere embryo B | 29.076 | 38.693 | 1.346 | 0.089 | WT/ altered paternal allele |
| MII | single blastomere embryo B | 29.208 | 35.472 | 1.850 | 0.132 | WT/ altered paternal allele |
| MII | single blastomere embryo B | 27.911 | 31.813 | 2.614 | 0.254 | WT/ altered paternal allele |
| MII | single blastomere embryo C | 29.076 | 38.693 | 1.350 | 0.111 | WT/ altered paternal allele |
| MII | single blastomere embryo C | 26.075 | 31.753 | 2.747 | 0.328 | WT/ altered paternal allele |
| MII | single blastomere embryo C | 26.878 | 30.841 | 2.669 | 0.292 | WT/ altered paternal allele |
| MII | single blastomere embryo C | 25.460 | 31.457 | 2.792 | 0.454 | WT/ altered paternal allele |
| MII | single blastomere embryo C | 32.133 | 33.847 | 2.170 | 0.383 | WT/ altered paternal allele |
| MII | single blastomere embryo C | no signal | no signal | na | na | no wt and no fs allele |
| MII | single blastomere embryo C | no signal | no signal | na | na | no wt and no fs allele |
| MII | TE BIOPSY (embryo E) | 27.508 | 32.761 | 2.731 | 0.377 | WT/ altered paternal allele |
| MII | TE BIOPSY (embryo F) | 31.511 | 36.146 | 2.257 | 0.179 | WT/ altered paternal allele |
| MII | SECOND TE BIOPSY (embryo F) | 28.242 | 33.515 | 2.672 | 0.312 | WT/ altered paternal allele |
| MII | TE biopsy embryo 10 (NHEJ) | 28.333 | 31.439 | 2.588 | 0.422 | WT/ altered paternal allele |
| MII | TE embryo 4 matching eysES4 | 28.622 | 30.936 | 2.660 | 0.338 | WT/ altered paternal allele |
| MII | TE embryo 4 matching eysES4 (second biopsy) | 29.893 | 34.743 | 2.025 | 0.271 | WT/ altered paternal allele |
| MII | TE embryo 5 matching eysES5 | 33.530 | 37.131 | 1.725 | 0.275 | WT/ altered paternal allele |
| MII | TE embryo 5 matching eysES5 (second biopsy) | 33.374 | 36.556 | 2.073 | 0.328 | WT/ altered paternal allele |
| MII | TE embryo 8 matching eysES8 | 36.278 | 39.119 | 0.764 | 0.208 | WT/ altered paternal allele |
| MII | TE BIOPSY embryo D (NHEJ) matches eysES9 | 26.971 | 34.739 | 2.484 | 0.242 | WT/ altered paternal allele |
| MII | SECOND TE BIOPSY embryo D (NHEJ) matches eysES9 | 27.992 | 36.703 | 2.057 | 0.192 | WT/ altered paternal allele |
| **Embryonic stem cell lines after MII RNP injection** | | | | | | |
| MII | eysES4 single cell | 27.214 | 33.449 | 2.467 | 0.274 | WT/ altered paternal allele |
| MII | eysES4 single cell | 28.019 | 35.135 | 1.943 | 0.217 | WT/ altered paternal allele |
| MII | eysES4 cell line (NHEJ) | 27.964 | 31.651 | 2.374 | 0.258 | WT/ altered paternal allele |
| MII | eysES5 cell line (NHEJ) | 27.635 | 30.812 | 2.538 | 0.327 | WT/ altered paternal allele |
| MII | eysES6 cell line (no change) | 27.868 | 25.968 | 2.318 | 3.306 | Heterozygous Carrier |
| MII | eysES6 single cell (no change) | 27.006 | 25.848 | 2.162 | 2.994 | Heterozygous Carrier |
| MII | eysES8 (NHEJ) | 27.790 | 31.498 | 1.226 | 0.266 | WT/ altered paternal allele |
| **INJECTION of Cas9 RNP AT 2-CELL STAGE** | | | | | | |
| 2-cell | single blastomere embryo 1 | no signal | 22.180 | 0.035 | 0.065 | no wt, no fs allele |
| 2-cell | single blastomere embryo 1 | no signal | no signal | na | na | no wt, no fs allele |
| 2-cell | single blastomere embryo 1 | 28.571 | 33.002 | 2.640 | 0.368 | WT/ altered paternal allele |
| 2-cell | single blastomere embryo 2 | 28.695 | 33.361 | 2.586 | 0.263 | WT/ altered paternal allele |
| 2-cell | single blastomere embryo 2 | 29.170 | 33.382 | 2.579 | 0.266 | WT/ altered paternal allele |
| 2-cell | single blastomere embryo 2 | 26.692 | 31.520 | 2.762 | 0.409 | WT/ altered paternal allele |
| **Other events** | | | | | | |
| 2PN | single blastomere embryo after 2PN injection | 32.199 | 36.867 | 2.055 | 0.131 | WT/ altered paternal allele |
| 2PN | single sister blastomere embryo after 2PN injection | 28.921 | 29.968 | 2.447 | 0.472 | WT/ altered paternal allele |

| sequence | Product size, Annealing Temp. | purpose |
|---|---|---|
| 5’-acagtgtgagagatgcccag | 699bp, 60 ℃ | PCR rs1482455, rs1482454, rs9362339 |
| 5’-tcaggactccaactaagcatga | 699bp, 60 ℃ | PCR rs1482455, rs1482454, rs9362339 |
| 5’-ggaagagcaaagcatctcca | na | Nested PCR rs1482455, rs1482454, rs9362339 |
| 5’-catcagaagaaatacaggagcca | na | Sequencing rs1482455, rs1482454, rs9362339 |
| 5’-acctaatctcatttcttggaggt | 765bp, 60 ℃ | PCR rs34809101, rs6936438 |
| 5’-aggatagcaacttcaaagactga | 765bp, 60 ℃ | PCR rs34809101, rs6936438 |
| 5’-actggtcttcgagctctgag | na | Sequencing rs3480910, rs6936438 |
| 5’-agatgagaaagaacaagcgca | na | Sequencing rs3480910, rs6936438 |
| 5’-gccctggagttgtttgtatga | Primer 8-10; 233bp, 60 ℃ | Primer 8; On target deep sequencing of mutation site and rs66502009. Indicated in Fig. 1A |
| 5’-tgaattttctggtgctttgttg | Primer 10-8; 233bp, 60 ℃; Primer 10-3; 1135bp | Primer 10; On target deep sequencing of mutation site and rs66502009. Indicated in Fig. 1A |
| 5'-acaggagaatggcatgaacc | Primer 2-11; 1536bp | Primer 2: Amplification and sequencing of rs66502009, mutation site, rs12205397, rs4530841, rs796414940; indicated in Fig. 1A |
| 5’-caaataagcaaaaatgagaaatgaaa | Primer 3-11: 1296bp | Primer 3; Amplification and sequencing of rs66502009, mutation site, rs12205397, rs4530841. Alternative for Primer 2, |
| 5’-ctttccccactctggagctgt | Primer 3-11: 1296bp | Primer 11; Amplification and sequencing of rs66502009, mutation site, rs12205397, rs4530841. |
| 5`- aaaatgtccacacttggctgg | na | Primer 7; Sequencing of mutation site, rs66502009 |
| 5’- cgctaatgtatcggctgg | na | Primer 6; Sequencing of rs12205397, rs4530841 |
| 5’-ctttccccactctggagctgt | 567bp, 60deg.C | Amplification of mutation site and rs66502009; alternative |
| 5`- aaaatgtccacacttggctgg | 567bp, 60deg.C | Amplification of mutation site and rs66502009; alternative |
| 5'-TGTACAAAAAAGCAGGCTTTAAAGGAACCAATTCAGTCGACTGGATCCGGTACCAAGGTCGGGCAGGAAGAGGGCCTATTTCCCATGATTCCTTCATATTTGCATATACGATACAAGGCTGTTAGAGAGATAATTAGAATTAATTTGACTGTAAACACAAAGATATTAGTACAAAATACGTGACGTAGAAAGTAATAATTTCTTGGGTAGTTTGCAGTTTTAAAATTATGTTTTAAAATGGACTATCATATGCTTACCGTAACTTGAAAGTATTTCGATTTCTTGGCTTTATATATCTTGTGGAAAGGACGAAACACCGGTGTCTTTCTTCTGTACTGGTTTTAGAGCTAGAAATAGCAAGTTAAAATAAGGCTAGTCCGTTATCAACTTGAAAAAGTGGCACCGAGTCGGTGCTTTTTTTCTAGACCCAGCTTTCTTGTACAAAGTTGGCATTA | na | gBLOCK for use in pluripotent stem cells |
| 5'-GUGUGUCUUUCUUCUGUACUGGUUUUAGAGCUAGAAAUAGCAAGUUAAAAUAAGGCUAGUCCGUUAUCAACUUGAAAAAGUGGCACCGAGUCGGUGCUUUU-3’ | na | guide RNA |
| 5'-TAATGCCAACTTTGTACAAGAAAG | 455bp, 57deg.C | Primer to amplify gBlock fragment |
| 5'-TGTACAAAAAAGCAGGCTTTAAAG | 455bp, 57deg.C | Primer to amplify gBlock fragment |
| 5'-AGGCTTGTCTCCCAACTAGA | 386bp, 57deg.C | forward primer for off-target analysis (chr8:37934195); site: TGTCTCTTTCTCTGTACTGAGG |
| 5'-ACATAAATTACATCACTGAGGGC | 386bp, 57deg.C | reverse primer for off-target analysis (chr8:37934195) |
| 5'-CCAGCCTCTTCACATATTTCC | 369 bp, 58deg.C | forward primer for off-target analysis (chr12:128575048); site: CTGTGTCTTTCTTTGTAATGTGG |
| 5'-AGTGTTGGGATTATAGGCATGA | 369 bp, 58deg.C | reverse primer for off-target analysis (chr12:128575048) |
| 5'-CTCTCGGTAGCTCTGGTT | 378bp, 55deg,C | forward primer for off-target analysis (chr7:2413257); site: GTTTTCTTCTGTACTGTGG |
| 5'-ATATGTTTGGCCACGTTCCT | 378bp, 55deg,C | reverse primer for off-target analysis (chr7:2413257) |
